# Sub-endothelial platelet activation amplifies neutrophil transmigration within venular walls

**DOI:** 10.64898/2026.07.21.739803

**Authors:** Tugce Cimen, Marilia Fernandes Manchope, Luisa Klinkenberg, Vanessa Göb, Beatriz Hoffmann Salles Bianchini, Natalia Reglero-Real, Wolfgang Kastenmüller, Ronen Alon, Bernhard Nieswandt, David Stegner, Tamara Girbl-Huemer

**Affiliations:** Rudolf Virchow Center for Integrative and Translational Bioimaging, Julius-Maximilians-Universität Würzburg (JMU), Würzburg, Germany; Institute of Experimental Biomedicine, Chair I, University Hospital Würzburg, Germany; Department of Immunology, Parasitology, and General Pathology, Center of Biological Sciences, Londrina State University, Londrina, Brazil; Departamento de Biología Molecular, Instituto Universitario de Biología Molecular (IUBM) and Centro de Biología Molecular Severo Ochoa (CBM), Universidad Autónoma de Madrid, UAM-CSIC, Madrid, Spain; Institute of Systems Immunology, University of Würzburg, Julius-Maximilians-Universität Würzburg (JMU), Würzburg, Germany; Department of Immunology and Regenerative Biology, Weizmann Institute of Science, Rehovot, Israel

## Abstract

Neutrophil recruitment into inflamed tissues requires coordinated transmigration across the multilayered venular wall, yet how this process is regulated beyond the endothelium remains poorly understood. Here, we identify a previously unrecognized platelet-neutrophil circuit that controls this post-endothelial phase. Intravital microscopy of inflamed cremasteric venules showed that pioneer neutrophil transmigration enabled platelet entry into the sub-endothelial compartment of venular walls. There, platelets became activated and released CXCL7 in a GPVI- and GPIbα-dependent manner, generating spatially confined chemokine microdomains. These platelet-derived cues directed follower neutrophil migration and promoted their exit across the pericyte layer into the interstitial tissue. Collectively, these findings identify extraluminal platelets as spatial organizers of neutrophil trafficking and uncover a localized intramural feedback mechanism in which neutrophils amplify their own recruitment within venular walls.

**Graphical summary:** 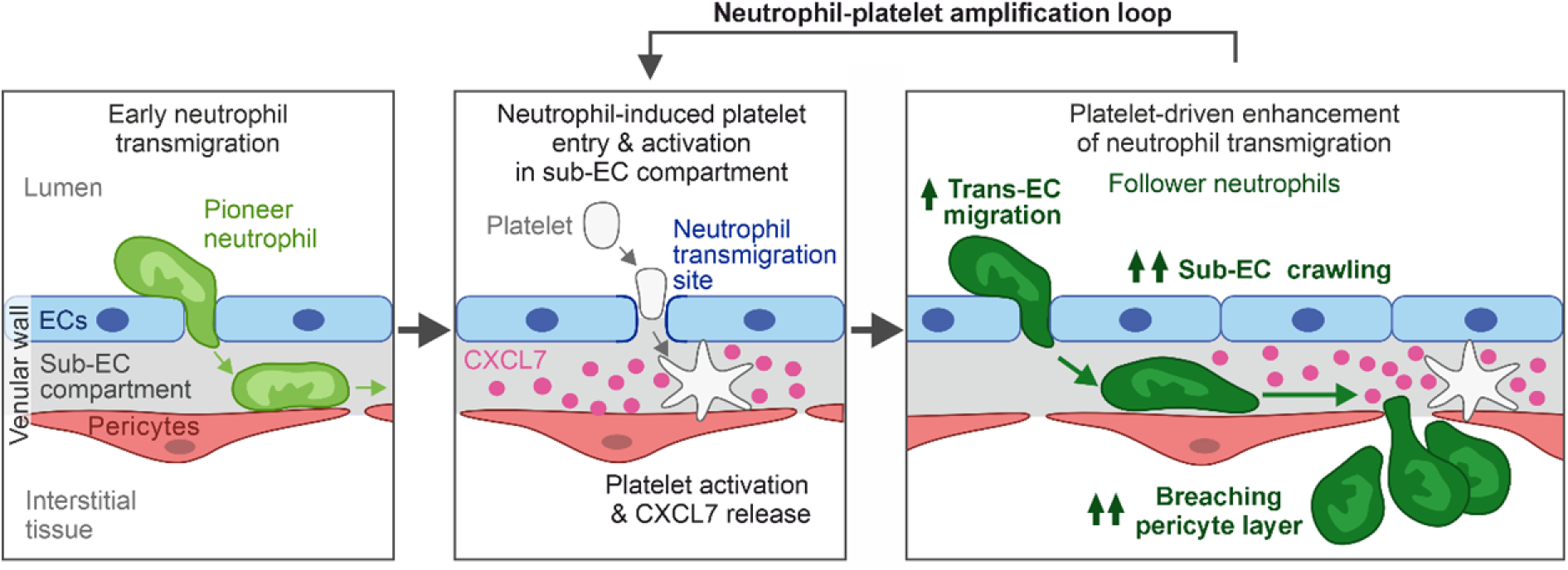

## Introduction

The recruitment of circulating neutrophils into inflamed tissues is a central component of host defense and a key driver of thrombo-inflammation. Neutrophils traverse inflamed venular walls through a complex, multistep cascade involving tethering, rolling, firm arrest, and intraluminal crawling on endothelial cells (ECs), followed by transendothelial migration and passage through the pericyte layer and associated venular basement membrane^1,2^. While the luminal phases of this cascade are well characterized and depend on coordinated interactions between neutrophils, ECs, and platelets^3–5^, how neutrophil behavior is spatially organized within the vessel wall after crossing the endothelium remains poorly defined.

After passing the endothelium, neutrophils extensively crawl along pericyte processes within the sub-endothelial (sub-EC) compartment of venular walls, before entering the inflamed interstitial tissue via gaps between adjacent pericytes^6^. This phase depends on pericyte-expressed ICAM-1 and CXCL1, prevents neutrophil reverse transendothelial migration back into the circulation, and is a prerequisite for neutrophil tissue entry^6,7^. Notably, neutrophil transmigration is spatially patterned and occurs preferentially at sites defined by endothelial function and pericyte architecture, basement membrane composition, and perivascular signals^6,8–11^. This spatial organization suggests the existence of local regulatory mechanisms within the vessel wall.

Platelets are key regulators of innate immunity and thrombo-inflammation^12,13^, and established modulators of luminal neutrophil behavior^5,14^. Through interactions involving GPIbα–von Willebrand factor (vWF), P-selectin–PSGL-1, integrins, and GPVI-dependent signaling, platelets promote neutrophil adhesion and intraluminal crawling^5,15–17^, and release soluble mediators such as serotonin metabolites, leukotriene B4, and chemokines that further enhance neutrophil recruitment^18–20^.

Platelets and platelet-derived structures, including microparticles and integrin- and tetraspanin-enriched tethers (PITTs), have been detected outside the vasculature during inflammation^21–23^, indicating that platelet functions extend beyond the circulation. The sub-EC compartment represents a matrix-rich microenvironment containing components such as collagen IV and laminins that can support platelet adhesion and activation^24^, raising the possibility that platelets acquire distinct functions upon entering this niche. However, whether platelets also impact neutrophil migration beyond the endothelial barrier remains unknown.

Here, using intravital microscopy (IVM) of inflamed cremaster muscle venules, we identify a previously unrecognized pathway in which pioneer neutrophil transmigration enables platelets to enter and accumulate within the sub-EC compartment of venular walls. Within this intramural niche, platelets become strongly activated and release CXCL7 in a GPVI and GPIbα-dependent manner, establishing spatially confined chemokine depots that guide follower neutrophils through the vessel wall and promote their exit across the pericyte layer.

Collectively, these findings define an intramural amplification mechanism in which pioneer neutrophils facilitate platelet entry into venular walls, and activated platelets in turn spatially organize follower neutrophil transmigration. This reveals that platelets actively coordinate immune cell trafficking from within the venular wall, thereby expanding their established role beyond the luminal endothelial interface.

## Results

### Individual platelets traverse EC junctions and are retained within the sub-EC compartment of venular walls during acute inflammation

To investigate platelet–neutrophil interactions *in vivo*, we induced acute inflammation in the murine cremaster muscle by intrascrotal (i.s.) tumor necrosis factor (TNF) administration (300 ng, 4 h)^6^, resulting in robust neutrophil recruitment (Figure 1A). Whole-mount immunofluorescence staining and confocal microscopy revealed that, in addition to luminal and junctional platelets (Figure S1A), TNF induced pronounced platelet entry and accumulation within the sub-EC compartment of postcapillary venules (Figures 1B and 1C; Video S1). Junctional platelets were defined as luminal platelets protruding into EC junctions, and sub-EC platelets were defined as having ≥50% of their volume beneath the endothelial layer.

Most sub-EC platelets (74.4 ± 2.0%, n = 24 vessels from 3 mice) were fully positioned between the endothelium and pericyte layer or embedded within pericyte processes (Figure 1B), while only a few retained small junctional protrusions. Nearly all directly contacted pericytes and the collagen IV-rich venular basement membrane (Figures 1B, S1B and S1C), whereas only few platelets were detected in the extravascular space (Figure 1C).

To define platelet entry dynamics, we utilized four-color confocal IVM and *Lyz2-eGFP-ki;Acta2-RFPcherry-Tg* reporter mice, which express eGFP^bright^ neutrophils and RFP^+^ pericytes^6^. EC junctions and platelets were labeled *in vivo* with non-blocking anti-CD31^25^ and anti-GPIX^26^ antibodies, enabling simultaneous high-resolution tracking of neutrophils and platelets relative to ECs and pericytes (Figures 1D–1F).

Neutrophils predominantly transmigrated paracellularly via EC junctions^25^ (Figure 1G). Strikingly, platelets adhered to the luminal endothelial surface near sites where neutrophils had completed transendothelial migration (lag time 3.8 min; Figures 1G and 1H) and subsequently crossed the endothelium through the same junctional site (Figure 1G; Video S2). Although neutrophil-induced EC junctional pores rapidly resealed, as reported^27^, transient discontinuities in CD31 staining frequently persisted and served as entry portals for platelets.

Platelet transendothelial passage was rapid (∼80% within 1 min; Figure 1I), compared to ∼6 min for neutrophils^7^. Overall, 92.3% of platelet entry events occurred at sites of prior neutrophil transendothelial migration (n = 7 mice). Despite frequently observing transient intraluminal platelet–neutrophil interactions, as previously described^28,16^, no joint transmigration was detected. Consistent with a causal link, neutrophil depletion (Figure S1D) markedly reduced platelet entry into the sub-EC compartment (Figure 1J).

Antibody-mediated blockade of the vWF-binding site of GPIbα strongly reduced sub-EC platelet accumulation, whereas GPVI immunodepletion from platelets had no major effect (Figure S1E). Neither intervention altered circulating platelet or polymorphonuclear leukocyte (PMN) counts (Figures S1F and S1G).

Sub-EC platelet accumulation was also observed in IL-1β- and LPS-induced cremasteric inflammation (Figures S1H–S1L) and during dermal inflammation after Modified Vaccinia Ankara virus (MVA) infection (Figures S1M–S1P), with neutrophil-dependence confirmed in LPS and MVA settings, indicating that this response is not restricted to cremasteric TNF stimulation.

Collectively, these data identify a neutrophil-enabled mechanism by which platelets traverse the endothelial barrier during inflammation, without requiring joint platelet-neutrophil transmigration. Notably, sub-EC platelet accumulation across multiple inflammatory settings indicates that this represents a recurring feature of acute inflammatory responses.

**Figure 1.**
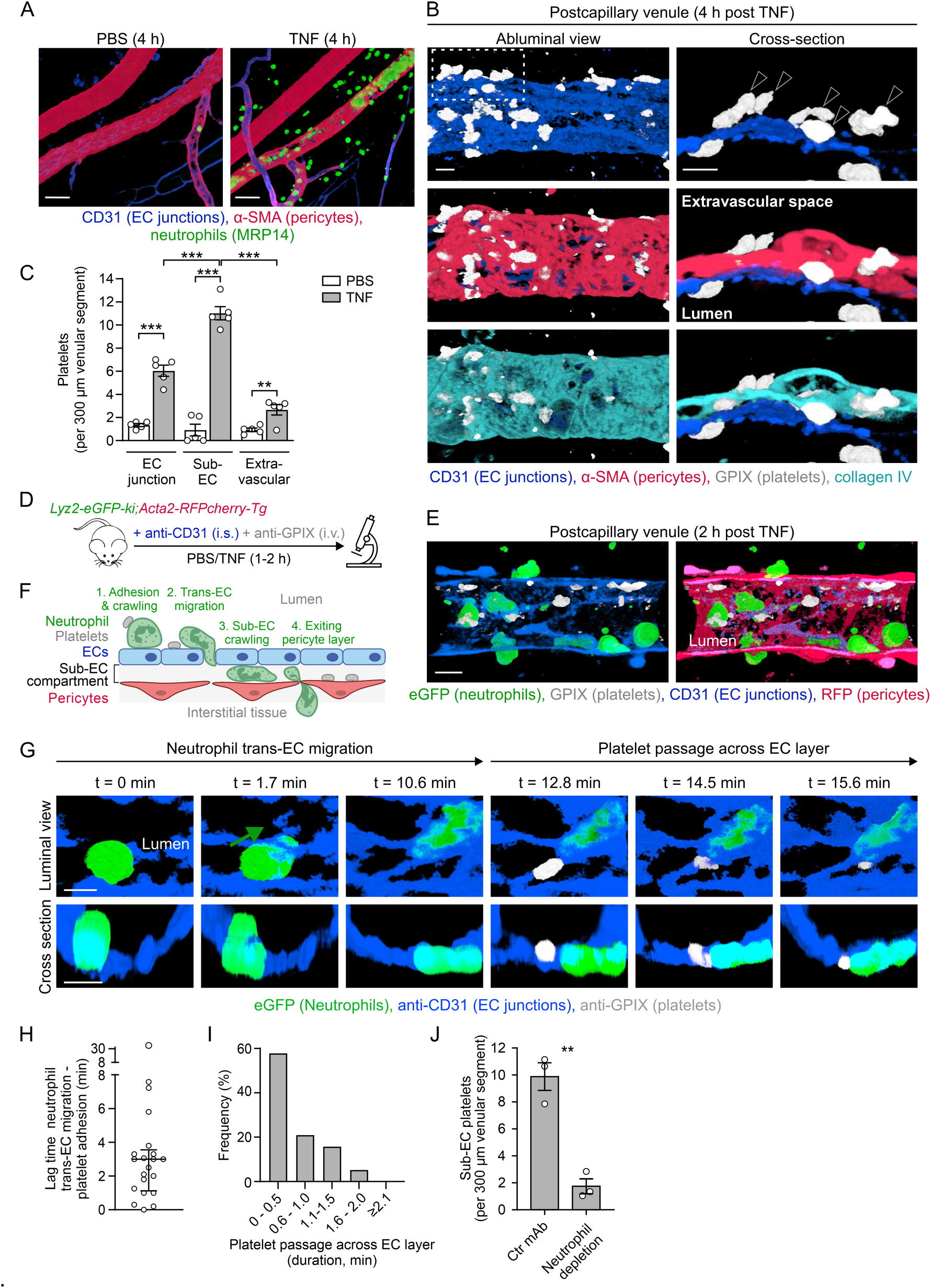
Individual platelets traverse EC junctions and are retained within the sub-EC compartment of venular walls during acute inflammation. **(A-C)** Wild-type (WT) mice received local (i.s.) injections of PBS or TNF. Whole-mount cremaster muscles were immunostained for CD31 (EC junctions), α-SMA (pericytes), MRP14 (neutrophils), GPIX (platelets), and/or collagen IV, and analyzed by confocal microscopy. **(A)** and **(B)** show representative images. Further examples of sub-EC platelets are shown in Video S1. Arrowheads indicate sub-EC platelets. Scale bars, 50 µm (A) and 5 µm (B). **(C)** Quantification of platelets at distinct localizations (n = 5 mice/group). **(D-I)** *Lyz2-eGFP-ki;Acta2-RFPcherry-Tg* mice were *in vivo* labeled with mAbs against CD31 and GPIX. Cremaster muscles were analyzed by confocal IVM, 2-4 h after i.s. TNF administration. **(D)** Scheme of confocal IVM procedures. **(E)** Representative projections of a 3D IVM image (scale bar, 10 µm). **(F)** Scheme of neutrophil and platelet responses investigated. **(G)** Time-lapse IVM images (Video S2) showing neutrophil adhesion (t = 0 min) and transendothelial migration (t = 1.7-10.6 min), followed by platelet adhesion (12.8 min) and passage across the EC junction (t = 14.5-15.6 min). For clarity, a representative neutrophil and platelet are shown, following isosurface rendering in Imaris. Scale bars, 5 µm. **(H)** Lag time between completion of neutrophil transendothelial (trans-EC) migration and platelet adhesion, and **(I)** duration of platelet translocation across EC junctions (n = 5 mice). **(J)** Sub-EC platelets in control or neutrophil-depleted (anti-Gr-1 mAb) mice (n = 3 mice/group), determined by confocal microscopy in cremaster muscles (4 h post TNF). Statistical significance was assessed by two-way ANOVA followed by Tukey’s post hoc test (C), or two-tailed Student’s *t* test with Welch’s correction (J). Means ± SEM, **p < 0.01, ***p < 0.001.

### Sub-EC platelets define neutrophil exit sites from the venular wall

Having established platelet accumulation within the sub-EC compartment, we next examined their behavior and impact on subsequent neutrophil transmigration.

During early inflammation (1–2 h after TNF), “pioneer” neutrophils – entering the sub-EC space before platelet arrival – slowly crawled along pericyte processes^6,7^ and exited the venular wall (Figure 2A). In contrast, sub-EC platelets remained largely stationary (mean displacement 2.0 ± 0.2 µm during 30 min; n = 7 mice) and did not enter the interstitial tissue (Figure 2A).

We next analyzed “follower” neutrophils entering venular segments already containing sub-EC platelets (2–4 h after TNF). These neutrophils crossed endothelial junctions at variable distances from platelets (Figures 2B and 2C), but once within the sub-EC compartment, preferentially migrated toward sub-EC platelets and exited the pericyte layer in their immediate vicinity (Figures 2B–2D, Video S3).

This spatial coupling was associated with frequent neutrophil–platelet contacts within the sub-EC compartment, where each platelet interacted with 4.0 ± 0.5 neutrophils with a mean contact duration of 9.5 ± 1.9 min (n = 4 mice). Notably, multiple neutrophils often exited sequentially through the same site in the pericyte layer, in close proximity to a sub-EC platelet (Figure 2E), indicating the formation of localized transmigration “hotspots”.

Furthermore, follower neutrophils exhibited increased sub-EC crawling speed and directional persistence compared to pioneers (Figures 2F and 2G) and a greater proportion exited the venular wall within 15 min (54.3% versus 6.3%, n = 5-8 mice/group), indicating enhanced migration dynamics associated with sub-EC platelet accumulation.

Collectively, these data demonstrate that platelets are stably retained within inflamed venular walls and mark preferential sites of directional neutrophil migration and exit through the pericyte layer.

**Figure 2.**
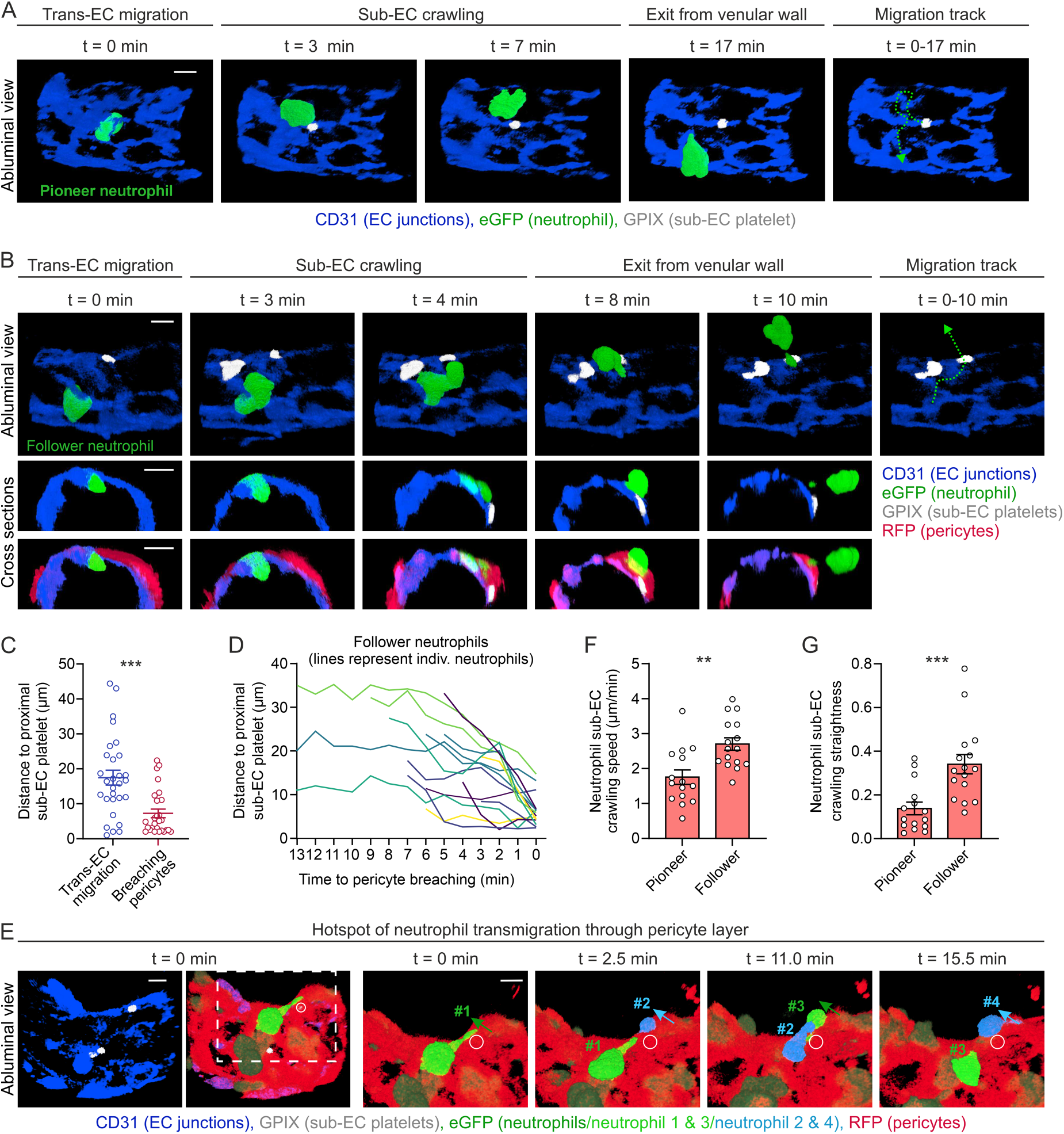
Sub-EC platelets mark preferential neutrophil exit sites from the venular wall. *Lyz2-eGFP-ki;Acta2-RFPcherry-Tg* mice were stained *in vivo* for CD31 and GPIX, i.s. injected with TNF, and cremasteric venules were analyzed by confocal IVM. **(A-B)** Time-lapse IVM images showing a **(A)** “pioneer” neutrophil during transendothelial migration (t =0 min), sub-EC crawling (t = 3-7 min), and exit from the venular wall (t = 17 min). A platelet enters the sub-EC compartment (t = 3 min), where it remains largely stationary (t = 3-17 min). Videos were recorded 1-2 h after TNF injection. Representative of 7 mice. **(B)** Images (Video S3) show a “follower” neutrophil during transendothelial migration (t = 0 min), sub-EC crawling (t = 3-4 min), and exit from the venular wall (t = 8-10 min) in relation to its proximal sub-EC platelet. Videos were recorded 2-4 h after TNF injection. Dashed lines indicate neutrophil sub-EC crawling tracks (n = 4 mice/group). Scale bars, 5 µm. **(C)** Distances between neutrophils and their proximal sub-EC platelet during neutrophil transendothelial migration and neutrophil breaching the pericyte layer (n = 29 and 27 neutrophils pooled from 4 mice/group). **(D)** Distances of individual sub-EC neutrophils to their proximal sub-EC platelet over time (n = 4 mice). **(E)** Neutrophil transmigration “hotspot”, showing four neutrophils migrating sequentially through the same area in the pericyte sheath close to a sub-EC platelet (circle). Images on the right show boxed area indicated in the overview. Representative of 5 mice. Scale bars, 5 µm. **(F-G)** Pioneer and follower neutrophil sub-EC crawling dynamics (displacement/track length, n = 15 and 16 neutrophils pooled from 3-4 mice/group). For clarity, sub-EC platelets and isolated neutrophils are shown/highlighted in (A, B, and E), following isosurface rendering in Imaris. Statistical significance was assessed by two-tailed Mann-Whitney U test (C and F) or two-tailed Student’s *t* test with Welch’s correction (G). Means ± SEM, **p < 0.01, ***p < 0.001.

### Platelets promote neutrophil sub-EC crawling and exit from venular walls

To directly assess the role of platelets in post-luminal neutrophil transmigration, we analyzed neutrophil passage across venular walls during TNF-induced inflammation after platelet depletion (Figure S2A). This protocol did not affect circulating PMN numbers (Figure S2B).

Platelet depletion reduced neutrophil adhesion and intraluminal crawling (Figures S2C and S2D), as reported^5,16^, as well as transendothelial migration (−36.1%; Figures 3A and 3B). Despite this partial inhibition, the majority of neutrophils (63.9%) still entered the sub-EC compartment, allowing further analyses (Figure 3C). In control mice, neutrophils displayed highly dynamic sub-EC crawling behavior, whereas platelet depletion markedly impaired this phase, reducing crawling speed and directional persistence (Figures 3D and 3E). This was accompanied by a decreased fraction of sub-EC neutrophils crossing the pericyte layer and increased retention of sub-EC neutrophils within venular walls (Figure 3F), consistent with the requirement of efficient sub-EC crawling for successful exit from venular walls^6,7^. The combined defects in luminal and sub-EC migration resulted in an overall 63.4% reduction in extravascular neutrophil accumulation (Figure 3G).

Overall, these results demonstrate that platelets regulate neutrophil transmigration beyond the luminal phase by promoting neutrophil sub-EC crawling and exit across the pericyte layer.

**Figure 3.**
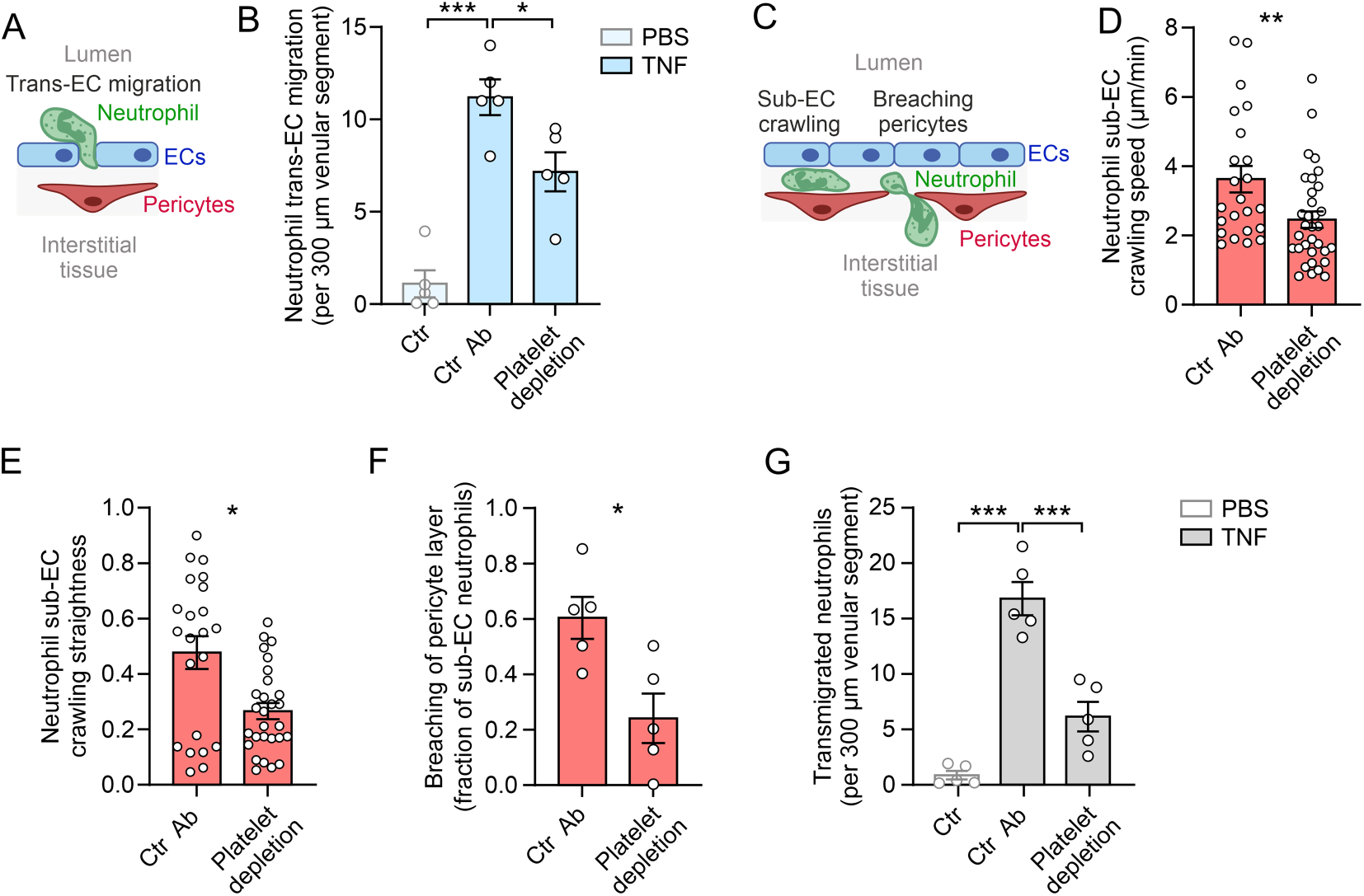
Platelets promote neutrophil sub-EC crawling and exit from venular walls. *Lyz2-eGFP-ki;Acta2-RFPcherry-Tg* mice were labeled *in vivo* for CD31 and GPIX and treated with control or platelet depleting anti-GPIbα antibodies (R300). Cremasteric venules were analyzed by confocal IVM 2-4 h after i.s. TNF administration. **(A)** Schematic illustrating neutrophil transendothelial migration quantified in (B). **(B)** Quantification of neutrophil transendothelial migration (n = 5 mice/group). Schematic of the neutrophil responses quantified in (D-F). **(D)** Neutrophil sub-EC crawling speed (n = 23 and 33 neutrophils pooled from 5 mice/group), **(E)** straightness index (n = 22 and 26 neutrophils from 5 mice/group), and **(F)** fraction of sub-EC neutrophils breaching the pericyte layer within 15 minutes (n = 5 mice/group). **(G)** Total transmigrated neutrophils in the interstitial tissue 4 h after PBS/TNF administration (n = 5 mice/group). Statistical significance was assessed using one-way ANOVA followed by Dunnett’s post hoc test (B and G), two-tailed Mann-Whitney U test (D and E), or unpaired two-tailed Student’s *t* test with Welch’s correction (F). Means ± SEM, *p < 0.05, **p < 0.01, ***p < 0.001.

### The sub-EC compartment induces robust platelet activation and α-granule chemokine release

Given that platelets cross the endothelium and promote neutrophil migration, we next examined whether the sub-EC compartment induces platelet activation. Because sub-EC access exposes platelets to extracellular matrix components that are normally concealed beneath the intact endothelium, such as collagen IV (Figure 1B; Figure S1C), we analyzed platelet calcium signaling using *Pf4-Cre;PC::G5-tdT* mice expressing the GCaMP5G calcium indicator and tdTomato specifically in megakaryocytes and platelets^29^ (Figure 4A). Platelet function in these mice was comparable to that of WT controls (Figures S3A-D; Table S1).

Confocal IVM showed low to intermediate calcium signals in luminal platelets, whereas sub-EC platelets exhibited markedly elevated GCaMP5G intensities (Figures 4B and 4C; Video S4), indicating enhanced activation. TdTomato signals, used as an internal fluorescence control, remained largely stable as expected (Figures S4A and S4B).

Tracking individual platelets revealed a rapid 3–4-fold increase in calcium upon entry into the sub-EC compartment (Figure 4D; Video S5), characterized by steadily increasing oscillatory calcium activity (platelet 1) or high-amplitude calcium spikes (platelets 2 and 3).

These findings indicate that translocation into the sub-EC compartment triggers rapid and robust platelet activation. To assess downstream effector functions, we examined platelet degranulation, through which activated platelets release mediators that modulate inflammatory responses^30^. Dense granule secretion did not measurably contribute to neutrophil extravasation, as *Unc13d^⁻/⁻^* mice, which lack platelet-dense granule secretion^31^, showed normal neutrophil extravasation (Figure S4C). We therefore focused on α-granule-derived mediators.

Platelet α-granules contain numerous immunomodulatory molecules, including adhesion receptors, cytokines, and chemokines. Among these, the *Cxcl7* gene products – platelet basic protein (PBP) and connective tissue-activating peptide III (CTAP-III) – constitute the most abundant platelet-derived chemokines and serve as precursors of NAP-2 (CXCL7). Myeloid-derived proteases convert these precursors into NAP-2, the potently chemotactic form that signals through CXCR1/CXCR2^32–34^. Throughout the study, we refer to NAP-2 and its precursors collectively as “CXCL7”.

Confocal microscopy showed that CXCL7 was almost exclusively platelet-associated in the cremaster microcirculation (Figure 4E), consistent with its dominant expression by megakaryocytes/platelets^34,35^. Nearly all luminal platelets were CXCL7⁺ under control and TNF conditions (Figures 4E–4H), and junctional platelets displayed similar levels (Figures 4G and 4H). In contrast, CXCL7 was undetectable or strongly reduced in sub-EC platelets (Figures 4G and 4H), consistent with substantial release upon entry into the sub-EC compartment. This spatial loss coincided with increased calcium signaling. Analysis of the α-granule chemokine CXCL4 (PF4) yielded similar results, with strong staining in luminal platelets and markedly reduced levels in sub-EC platelets (Figures S4D–S4E).

Furthermore, ELISA of whole cremaster tissue (including intravascular blood) showed comparable CXCL7 levels under control and TNF conditions, whereas platelet depletion reduced CXCL7 by ∼93% (Figure 4I), identifying platelets as the predominant source.

Collectively, these data identify the sub-EC compartment as a specialized site of robust platelet activation and α-granule release, characterized by calcium signaling and the release of CXCL7 and CXCL4.

**Figure 4.**
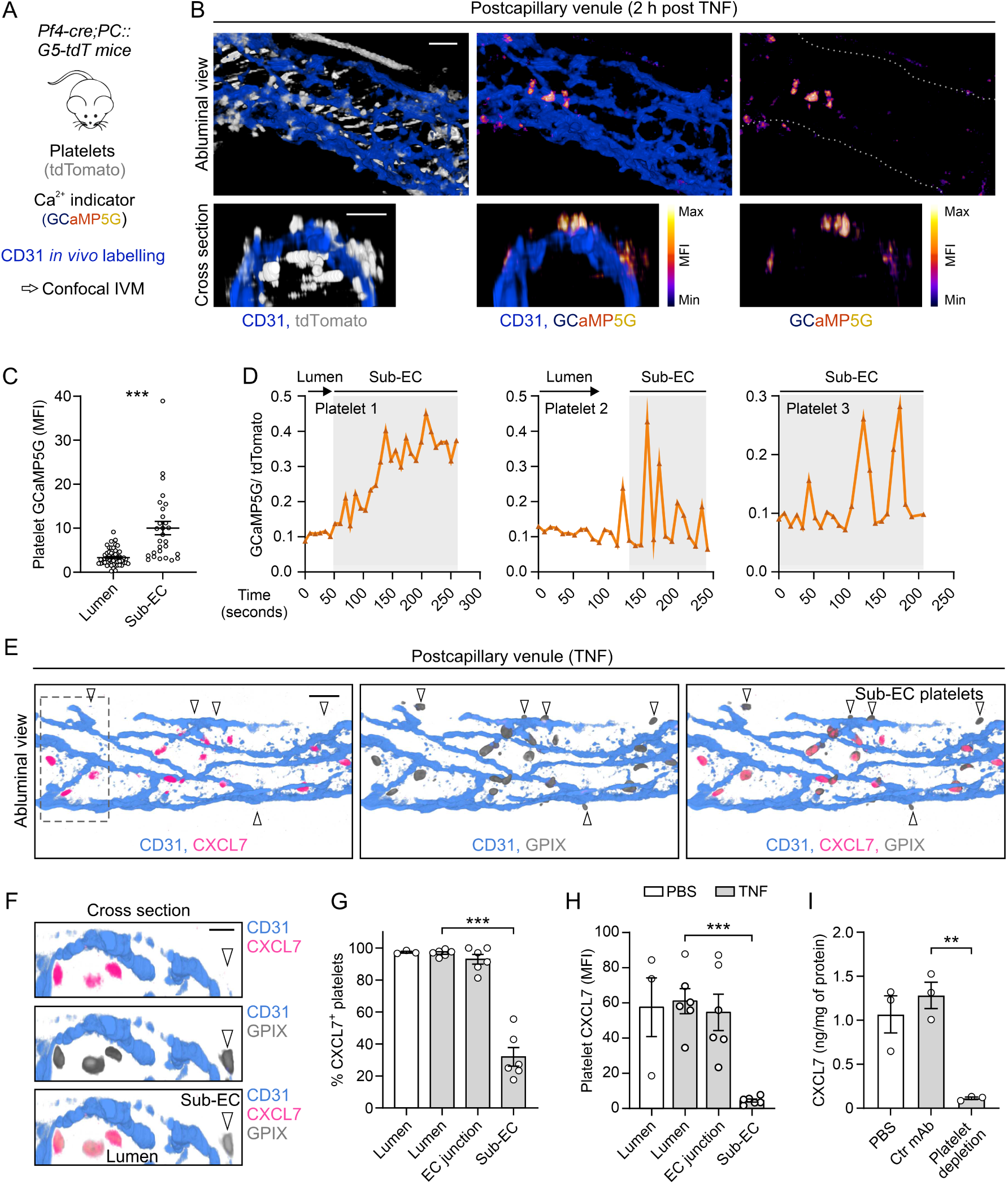
The sub-EC compartment induces robust platelet activation and α-granule chemokine release. (A-D) Platelet calcium signaling was analyzed by confocal IVM in *Pf4-Cre;PC::G5-tdT* mice, which express tdTomato and the calcium indicator GCaMP5G, specifically in megakaryocytes and platelets, 2-4 h after i.s. TNF administration. Endothelial junctions were labeled *in vivo* with anti-CD31 antibodies **(A)**. **(B)** Representative IVM images. Scale bars, 10 µm. **(C)** GCaMP5G (GFP) MFIs in luminal and sub-EC platelets in TNF-stimulated cremasteric venules (n = 51 and 28 platelets pooled from 3 mice/group). Representative GCaMP5G (GFP)/tdTomato fluorescence traces from individual platelets during their translocation from the lumen into the sub-EC compartment or after entry into this compartment. **(E-H)** Cremaster muscles from WT mice were *in vivo* stained for CD31, collected 4 h after PBS or TNF administration, immunostained for GPIX and CXCL7, and analyzed by confocal microscopy. **(E)** Representative overview and **(F)** cross-sections of the boxed region in (E). Arrowheads indicate sub-EC platelets. Scale bars, 10 µm. **(G-H)** Quantification of platelet-associated CXCL7 signals (n = 3-6 mice/group). **(I)** ELISA quantification of CXCL7 in cremaster muscles of WT mice after control or anti-GPIbα antibodies (R300) administration for platelet depletion, and PBS or TNF stimulation. (n = 3 mice/group). Statistical significance was assessed using two-tailed Mann-Whitney U test (C), or one-way ANOVA followed by Dunnett’s post hoc test (G-I). Means ± SEM, **p < 0.01, ***p < 0.001.

### GPVI and GPIbα mediate CXCL7 release from sub-EC platelets

Platelet adhesion receptors are central regulators of platelet activation in hemostatic and inflammatory settings. GPVI, the principal collagen receptor, and GPIbα, which mediates adhesion via vWF, have both been implicated in neutrophil recruitment^14,15,36,37^.

To assess their role in sub-EC platelet function, we performed GPVI immunodepletion and Fab-mediated GPIbα blockade (Figures S1E–S1G). Neither intervention altered CXCL7 levels in luminal platelets (Figures 5A–5C). GPVI depletion had no effect on junctional platelets, whereas GPIbα blockade modestly increased the fraction of CXCL7⁺ junctional platelets without affecting overall signal intensity (Figures S5A and S5B). In contrast, inhibition of either GPVI or GPIbα markedly increased CXCL7 retention in sub-EC platelets, consistent with impaired chemokine release (Figures 5D and 5E). Combined inhibition further enhanced this effect, with ∼70% CXCL7 signal retained compared to ∼7% under control conditions (Figure 5E).

We next examined the functional consequences of platelet receptor inhibition on neutrophil migration. Among neutrophils that had entered the sub-EC compartment, both GPVI immunodepletion and GPIbα blockade reduced neutrophil sub-EC crawling speed and directional persistence and decreased the fraction of sub-EC neutrophils crossing the pericyte layer (Figures 5F–5H, 5J–5L). Consistent with these findings, GPVI immunodepletion and GPIbα blockade, reduced the overall number of extravascular neutrophils (Figures 5I and 5M).

Because GPIbα blockade also impaired platelet entry into the sub-EC compartment (Figure S1E), its effects cannot be attributed solely to altered CXCL7 release. Nonetheless, these findings identify GPVI and GPIbα as key regulators of platelet effector function within the venular wall, promoting CXCL7 release and platelet-mediated neutrophil guidance through the sub-EC compartment. We therefore directly examined the role of CXCL7 in neutrophil trafficking.

**Figure 5.**
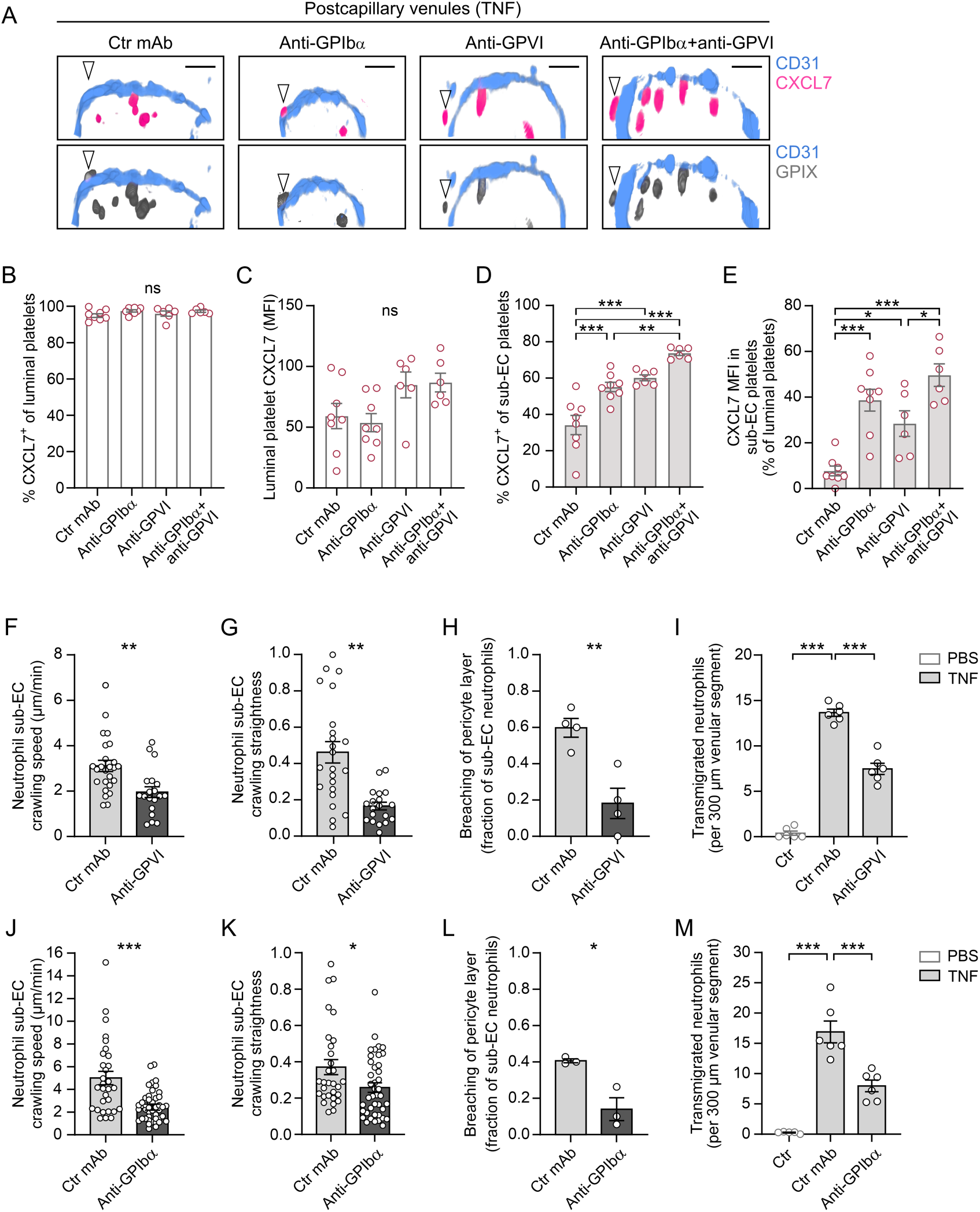
GPVI and GPIbα mediate CXCL7 release from sub-EC platelets. WT or *Lyz2-eGFP-ki;Acta2-RFPcherry-Tg* mice were treated with control Fabs or Fab fragments of an anti-GPIbα mAb and/or pretreated with control or anti-GPVI mAbs to immunodeplete GPVI from circulating platelets. **(A-E)** Cremasteric venules from WT mice were immunostained and analyzed by confocal microscopy 4 h after TNF administration. **(A)** Representative images. Arrowheads indicate sub-EC platelets. Scale bars, 10 µm. **(B-C)** Quantification of CXCL7 signals in luminal platelets, and **(D-E)** in sub-EC platelets. Results from the corresponding Fab and whole-IgG control groups were similar and therefore pooled (n = 6-8 mice/group). Signals in sub-EC platelets were normalized to luminal platelets (=100%) for each mouse in (E). **(F-H** and **J-L)** Confocal IVM analysis of cremasteric venules from *Lyz2-eGFP-ki;Acta2-RFPcherry-Tg* mice 2-4 hours after TNF administration. **(F, J)** Neutrophil sub-EC crawling speed (n = 25 and 19 (F), or 30 and 46 (J) neutrophils from 3-4 mice/group) and **(G, K)** straightness index (n = 23 and 19 (G), or 30 and 41 (K) neutrophils from 3-4 mice/group). **(H, L)** Fraction of sub-EC neutrophils breaching the pericyte layer (n = 4 mice/group in F-H, and n = 3 mice/group in J-L). **(I, M)** Quantification of extravascular neutrophils in WT mice subjected to indicated antibody treatments, determined by confocal microscopy 4 h after PBS/TNF administration. (n = 6 mice/group). Statistical significance was assessed using one-way ANOVA followed by Tukey’s post hoc test (B-E), unpaired two-tailed Student’s *t* test with Welch’s correction (F-H and L), one-way ANOVA followed by Dunnett’s post hoc test (I and M), or an unpaired two-tailed Mann-Whitney U test (J and K). Means ± SEM, *p < 0.05, **p < 0.01, ***p < 0.001, ns, not significant.

### CXCL7 drives neutrophil sub-EC crawling and exit from venular walls

Building on our finding that sub-EC platelets release CXCL7 and direct neutrophil migration, we next examined whether CXCL7 acts as an intramural guidance cue. Because TNF-induced neutrophil extravasation strongly depends on CXCR2^7^ and the potency of CXCL7 as a CXCR2 agonist, we tested whether CXCL7 functions as an intramural guidance cue within venular walls.

Intravenous (i.v.) administration of a blocking anti-CXCL7 mAb did not affect neutrophil adhesion or intraluminal crawling (Figures 6A and 6B), indicating that CXCL7 is dispensable for these luminal phases of transmigration.

To assess post-luminal effects, blocking antibodies were administered i.s. 110 min after TNF injection, as previously established^7^. Under these conditions, CXCL7 inhibition reduced neutrophil transendothelial migration (Figure 6C) and significantly impaired sub-EC crawling speed and directional persistence (Figures 6D–6F). Moreover, CXCL7 blockade significantly impaired sub-EC neutrophil passage across the pericyte layer, leading to increased retention within the venular wall (Figure 6G).

Consistent with an intramural guidance function, CXCL7 blockade disrupted neutrophil sub-EC crawling toward platelets (Figure 6H). Platelet numbers in both luminal and sub-EC compartments were unchanged (Figures S6A and S6B), supporting the conclusion that the migratory defects were not caused by altered platelet localization.

Together, these data identify CXCL7 as a key platelet-derived chemotactic cue that regulates neutrophil migration within the venular wall and promotes their exit across the pericyte layer during inflammation.

**Figure 6.**
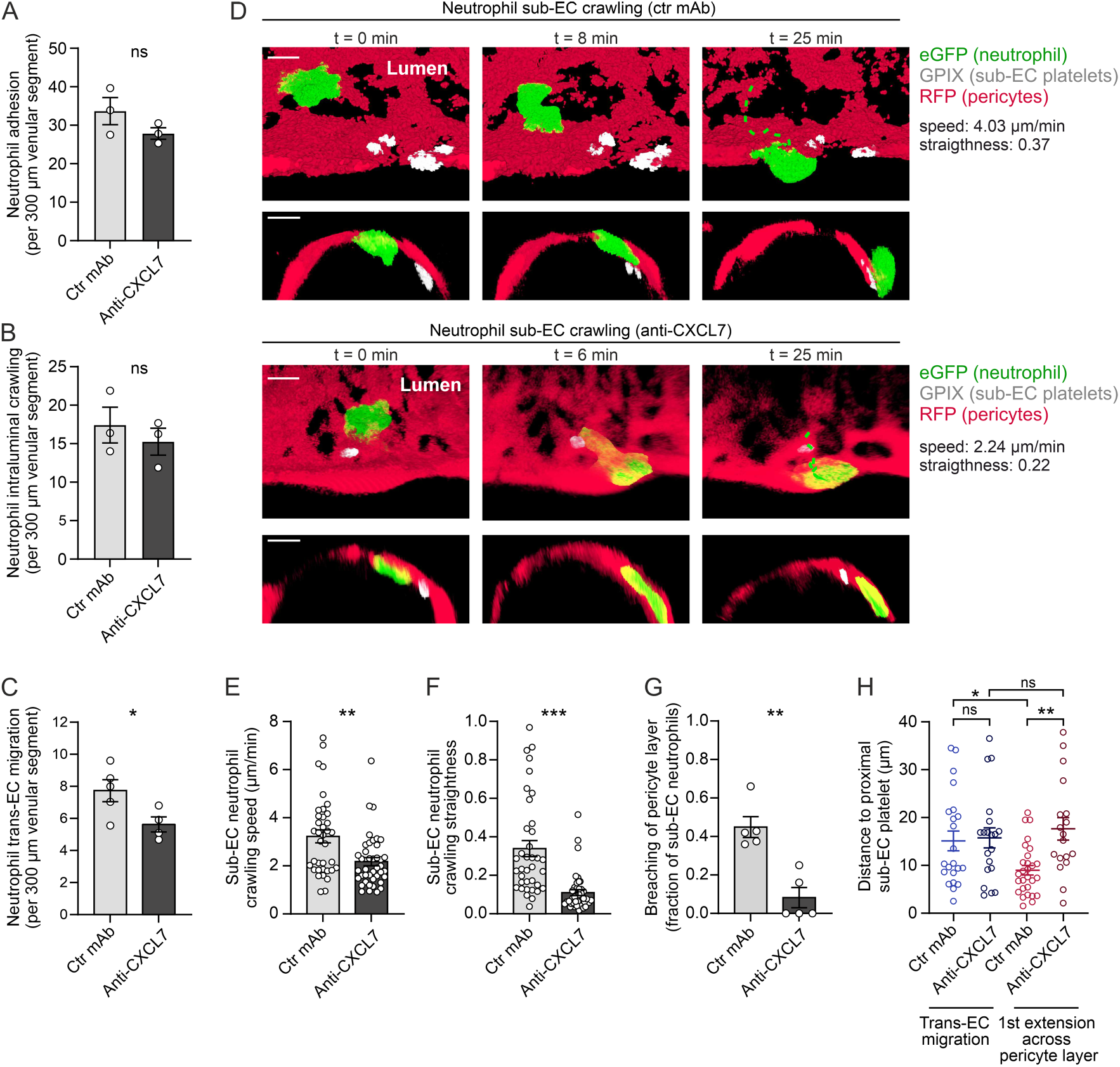
CXCL7 drives neutrophil sub-EC crawling and exit from venular walls. Cremasteric venules of *Lyz2-eGFP-ki;Acta2-RFPcherry-Tg* mice were analyzed by confocal IVM 2-4 h after TNF administration. Mice received control or blocking anti-CXCL7 mAbs i.v. 10 min before TNF administration for analyses in (A, B), or i.s. 110 min after TNF for analyses in (C-H) and were labeled *in vivo* for CD31 and GPIX. **(A-B)** Neutrophil adhesion and intraluminal crawling (n = 3 mice/group). **(C)** Neutrophil transendothelial migration (n = 4-5 mice/group). **(D)** Representative IVM images showing neutrophil sub-EC crawling along pericytes after control or anti-CXCL7 mAb treatment. Dashed lines indicate crawling tracks. For clarity, a selected neutrophil and sub-EC platelets are shown following isosurface rendering. Scale bars, 5 µm. **(E-F)** Neutrophil sub-EC crawling speed and straightness index (n = 37 and 41 neutrophils pooled from 5 mice/group), and **(G)** fraction of sub-EC neutrophils crossing the pericyte layer (n = 5 mice/group). **(H)** Distances between neutrophils and their nearest sub-EC platelets during neutrophil transendothelial migration and at the first protrusion across the pericyte layer (n = 18-28 neutrophils pooled from 3-5 mice/group). Statistical significance was assessed using unpaired two-tailed Student’s *t* test with Welch’s correction (A-C), or two-tailed Mann-Whitney U test (E-G), or two-way ANOVA (H). Means ± SEM, *p < 0.05, **p < 0.01, ***p < 0.001, ns, not significant.

### Platelets enter the sub-EC compartment during ischemia–reperfusion (I/R) injury

To test whether platelet–neutrophil interactions extend beyond TNF-induced inflammation, we analyzed an I/R injury model in the cremaster muscle^8^. 40 min of ischemia followed by 3 h of reperfusion induced robust neutrophil extravasation (Figures 7A and 7B).

Furthermore, I/R injury led to pronounced platelet accumulation within endothelial junctions and the sub-EC compartment of venular walls, but not in the extravascular space, while platelet numbers remained low in sham controls (Figures 7C and 7D). Neutrophil depletion (Figure S7A) markedly reduced platelet entry into junctional and sub-EC compartments (Figure 7D), indicating neutrophil dependence.

Local CXCL7 blockade reduced overall neutrophil transmigration during I/R injury (Figures 7A and 7B) without affecting platelet positioning (Figures S7B and S7C), supporting a direct role for platelet-derived CXCL7.

Together, these findings indicate that sub-EC platelet accumulation also occurs during I/R injury and support a role for platelet-derived CXCL7 in neutrophil extravasation in this pathological context.

**Figure 7.**
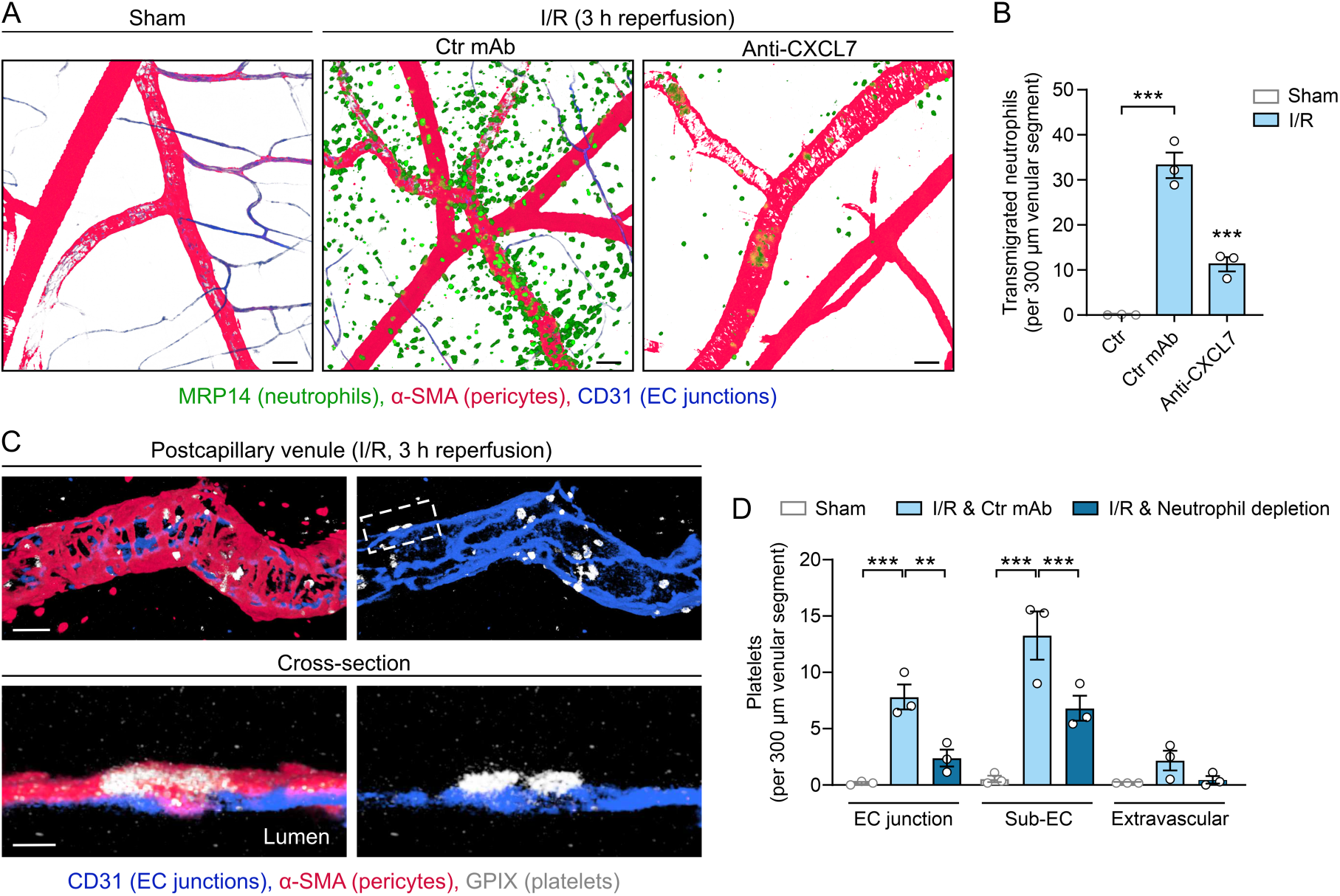
Platelets enter the sub-EC compartment during I/R injury. Sham or I/R injury of cremaster muscles of WT mice analyzed by confocal microscopy. **(A)** Representative images (scale bars, 50 µm) and **(B)** overall transmigrated neutrophils. Control or blocking anti-CXCL7 mAbs were administered i.s. (n = 3 mice/group). **(C)** Images showing two sub-EC platelets. Scale bars, top: 10 µm; bottom: 2 µm. **(D)** Platelet location in control or neutrophil-depleted (anti-Ly6G mAb) mice (n = 3 mice/group). One-way ANOVA followed by Dunnett’s post hoc test (B), or two-way ANOVA followed by Tukey’s post hoc test (D). Means ± SEM, **p < 0.01, ***p < 0.001.

## Discussion

Thrombo-inflammation couples platelet activation to immune cell recruitment and tissue injury, yet how these processes are coordinated within the vessel wall has remained unclear. While neutrophil recruitment has classically been viewed to be regulated at the luminal endothelial surface, our findings extend this paradigm by identifying a platelet-dependent mechanism operating within the sub-EC compartment that shapes neutrophil trafficking after transendothelial migration. In this context, platelets enter the vessel wall and establish localized chemokine microdomains that guide neutrophil exit into the tissue. This creates a spatially confined amplification circuit in which pioneer neutrophil transmigration enables platelet entry, which in turn promotes the passage of follower neutrophils across the vessel wall. Together, these findings define an additional spatial layer of regulation in immune cell recruitment. Platelets are established mediators of luminal neutrophil migration via P-selectin– PSGL-1 and integrin-dependent interactions^5,14^. Our data now place platelet functions within a distinct intramural context, where platelets act as local organizers of neutrophil trafficking, extending their established functions at the luminal endothelial interface to spatial control of leukocyte migration within the venular wall.

Mechanistically, platelets access the sub-EC compartment by traversing transient junctional discontinuities that persist after neutrophil diapedesis, without evidence of joint transmigration, arguing against a “piggy-back” mechanism. While platelet accumulation at sites of leukocyte extravasation and within endothelial junctions has been reported^17,21^, our data demonstrate that platelets fully cross the endothelium and are retained between endothelial and pericyte layers, establishing a distinct intramural compartment. This localization places platelets in a microenvironment that is shielded from luminal flow, enabling spatially restricted signaling.

Beyond regulating leukocyte recruitment, platelets are well established to maintain vascular integrity during inflammation^38^. Previous work showed that platelets seal diapedesis-associated breaches and prevent leakage via Angiopoietin-1–Tie2 signaling^21,39,40^. Our findings extend this model by showing that platelets do not only enter EC junctions but frequently fully traverse the endothelium, where they actively promote neutrophil trafficking via localized chemokine release. Thus, platelet recruitment to venular walls integrates barrier protection with control of leukocyte migration at successive stages of diapedesis.

Platelet entry into the sub-EC compartment depended primarily on GPIbα, consistent with its role in vWF-mediated adhesion and neutrophil extravasation downstream of leukocyte adhesion^43,47^. In contrast, GPVI immunodepletion did not alter platelet entry despite prior reports implicating GPVI in platelet accumulation at diapedesis sites^39^, suggesting redundancy with alternative adhesion pathways^21^. However, GPVI was required for CXCL7 release, consistent with platelet activation by GPVI ligands exposed within the sub-EC extracellular matrix.

This provides a mechanistic explanation for the importance of GPVI in thrombo-inflammation^42^: in the settings studied here, GPVI was dispensable for platelet entry, but required for intramural effector function. Consistent with this concept, partial GPVI downregulation was recently shown to protect against LPS-induced thrombo-inflammation while preserving hemostasis^43^, suggesting that selective attenuation of GPVI-dependent platelet effector functions may offer a therapeutically favorable strategy. Together with our identification of platelet-derived tethers (PITTs) as regulators of thrombo-inflammation^23^, this supports a model in which platelet adhesion receptors orchestrate immune responses within spatially defined niches.

Among α-granule mediators, CXCL7 emerged as a key effector. CXCL7 has been implicated in neutrophil guidance within thrombi and in tissue regeneration^35,44^. Here, we identify the sub-EC compartment as a site of CXCL7 release. In contrast to luminal chemokines, which are subject to rapid dilution by blood flow, chemokines released in the sub-EC space are protected from luminal dilution and likely stabilized by glycosaminoglycans^45–47^. Under these conditions, CXCL7 establishes spatially confined intramural guidance cues directing neutrophils toward platelet-rich regions and permissive sites of exit across the pericyte layer.

How platelet-derived CXCL7 facilitates not only neutrophil sub-EC crawling, but also neutrophil passage across the pericyte layer remains unresolved. CXCL7 may locally modulate CXCR2 responsiveness or act in concert with platelet-mediated changes in pericytes or the basement membrane, but these possibilities require direct experimental investigation.

Similar self-reinforcing amplification loops have been described within interstitial tissues, where neutrophil-derived signals mediate collective migration^48,49^, suggesting that such feedback mechanisms may represent a general principle of immune cell recruitment across tissue compartments.

Our previous work demonstrated that spatially compartmentalized chemotactic cues, including CXCL1 and CXCL2, exhibit distinct and non-redundant functions in neutrophil transmigration through signaling via CXCR2^7^. In this context, CXCL1, derived from ECs and pericytes, promotes neutrophil luminal and sub-EC crawling, whereas CXCL2, primarily produced by neutrophils, regulates transendothelial migration. Our current data extend these findings by showing that platelet-established CXCL7 microdomains provide neutrophil guidance cues that are essential for their efficient tissue entry. These findings suggest that CXCL7 acts in concert with CXCL1 to coordinate efficient neutrophil transmigration beyond the endothelium. While pericyte-derived CXCL1 is broadly distributed and supports basal non-directional sub-EC crawling^7^, platelet-derived CXCL7 provides spatially confined chemotactic microdomains that direct neutrophils towards platelet-rich regions and permissive exit sites in the pericyte layer.

Consistent with this function, we frequently observed multiple neutrophils sequentially exiting through the same regions of the pericyte layer closely aligned with sub-EC platelets. These findings suggest that platelet-derived cues help establish or reinforce neutrophil transmigration hotspots within the pericyte layer. We have previously shown that mast cell–dependent pericyte activation can contribute to the formation of such sites^11^, suggesting that platelets and mast cells might act through complementary pathways to localize neutrophil egress from vessel walls.

Finally, we demonstrate that sub-EC platelet accumulation and CXCL7-dependent neutrophil recruitment also occur during I/R injury, indicating that this mechanism is not restricted to TNF-induced inflammation. These observations suggest that platelet-mediated intramural guidance may represent a general feature of inflammatory responses across diverse pathological contexts.

In summary, we identify a previously unrecognized intramural platelet function that amplifies inflammatory neutrophil recruitment. By entering the venular wall and establishing localized chemokine microdomains, platelets act as spatial organizers and amplifiers of neutrophil recruitment. This mechanism may help explain the prominent roles of platelet receptors, including GPIbα and GPVI, in thrombo-inflammation, by enabling platelet function within protected extraluminal compartments that directly control immune cell behavior.

These findings provide new insights into how immune cell recruitment is coordinated across distinct vascular compartments and highlight intramural signaling as a key layer of regulation in inflammatory responses.

## Acknowledgments

We thank Dr. Sussan Nourshargh and Dr. David Rowe for providing the *Acta2-RFPcherry-Tg* mice and the Core Unit Fluorescence Imaging of the Rudolf Virchow Center for their valuable technical support. We thank Dr. Michaela Kuhn and the machine shop of the Institute of Physiology at the University of Würzburg for fabrication of customized microscope stages and the undergraduate students Kira Weitner, Isabella Bergner, and Aina Freixedas Raventós for their assistance. This work was supported by institutional funding from the Rudolf Virchow Center, University of Würzburg and the Deutsche Forschungsgemeinschaft (DFG, German Research Foundation; RTG 3190, project number 556462806; CRC1525, project number 453989101). B.H.S.B. was supported by the PhD scholarship from the Coordination for the Improvement of Higher Education Personnel (CAPES, Brazil, finance code 001).

## Author contributions

Conceptualization and writing (original draft): B.N., D.S., M.F.M., T.C., T.G.H.; Funding acquisition: B.N., D.S., T.G.H.; Methodology: B.N., D.S., M.F.M., T.C., T.G.H., V.G., W.K.; Investigation and formal analysis: B.H.S.B., L.K., M.F.M., N.R.R., T.C., T.G.H., V.G.; Resources: B.N., D.S., T.G.H., W.K.; Supervision: B.N., D.S., R.A., T.G.H.; Writing (review and editing): All authors.

## Methods

### Mice

WT C57BL/6J mice were obtained from Charles River (Sulzfeld, Germany). *Lyz2-eGFP-ki* mice were kindly provided by Dr. Thomas Graf (Center for Genomic Regulation and ICREA, Spain). These mice express eGFP^+^ myeloid cells (eGFP^bright^ neutrophils and eGFP^dim^ monocytes) by knocking an eGFP gene into the lysozyme M (*Lyz2*) locus^50^. *Acta2-RFPcherry-Tg* mice were kindly provided by Dr. Sussan Nourshargh (Center for Microvascular Research, WHRI, Queen Mary University of London, UK) and used with permission of Dr. David Rowe (Center for Regenerative Medicine and Skeletal Development, University of Connecticut, US). This colony exhibits a transgenic insertion of the RFP variant mCherry under the control of *Acta2* promoter and expresses RFP^+^ pericytes and smooth muscle cells^6^. *Lyz2-eGFP-ki;Acta2-RFPcherry-Tg* mice were generated by crossing the *Lyz2-eGFP-ki* colony with the *Acta2-RFPcherry-Tg* colony. *Unc13d*^-/-^ mice were generated by Velocigene and have a point mutation in the *Unc13d* gene, leading to the absence of Munc13-4 in platelets, abolishing the release of dense granules from platelets^31^. *Pf4-Cre* mice^51^ were generously provided by Dr. Radek C. Skoda (Department of Biomedicine, University of Basel, Switzerland). *Pf4-Cre;PC::G5-tdT* mice were generated by crossing *PC::G5-tdT* mouse line^29^ with a *Pf4-Cre* mouse line. The progeny have platelets expressing the fluorescent calcium sensor variant GCaMP5G and tdTomato fluorescent protein under the control of the CAG promoter targeted in an intergenic region downstream of the endogenous *Polr2a* gene. All mice were housed in individually ventilated cages under specific pathogen-free (SPF) conditions and a 12 h light-dark cycle at the Rudolf Virchow Center or the Center for Experimental Molecular Medicine at the University of Würzburg or the Universidad Autónoma de Madrid. Room temperature was maintained at 22-24°C and humidity was maintained at 50-60%. Food and water were provided *ad libitum*. Male mice were used to investigate inflammatory responses in the cremaster muscle, and females were used to study dermal inflammation. All experiments were conducted using 8-to 12-week-old mice that were age-matched. Mouse experiments were approved by the district government of Lower Franconia (Bezirksregierung Unterfranken) and the Spanish institutional review boards (Centro de Biología Molecular Severo Ochoa, Consejo Superior de Investigaciones Científicas and Madrid Community Ethical Committee, Spain).

### Inflammatory responses in the cremaster muscle

Mice were anesthetized with 3% isoflurane and received i.s. injections of 300 ng TNF (R&D Systems), 500 ng of LPS (Sigma-Aldrich), or 50 ng of IL-1β (BioLegend) in 350 µL PBS (Thermo Fisher Scientific), whereas control mice were injected with only PBS and were incubated for 1-4 h. I/R injury was induced in cremaster muscles by placing two orthodontic elastic bands (1/8”, 4.5 Oz) around the proximal testicular region for 40 min. The elastic bands were removed, and cremaster muscles were re-perfused for 3 h^8^.

### Depletion of platelets or neutrophils

For platelet depletion, mice were treated i.v. with anti-GPIbα antibodies (R300, Emfret Analytics), or an isotype-matched control (rat IgG, C301, Emfret Analytics) antibody^52^ at a dose of 2 µg/g body weight, 14 h before induction of cremasteric inflammation. This led to an >95% reduction in circulating platelet counts, as evaluated with an automated hemocytometer at the end of each experiment. Additionally, the cumulative volume of platelets tethered within the lumen of cremasteric venular segments was analyzed by immunofluorescence staining and confocal microscopy and defined as platelets retained within the lumen after tissue fixation, washing and staining. Depletion of circulating neutrophils was performed by intraperitoneal (i.p.) injections of an anti-Gr-1 mAb (clone RB6-8C5, BioLegend, 25 µg/day for 3 days^53^). The last dose was administered 1 h before i.s. TNF or LPS injection. Alternatively, circulating neutrophils were depleted by i.p. injections of an anti-Ly6G mAb (clone 1A8, BioLegend, 25 µg/day) together with a secondary anti-Kappa Immunoglobulin Light Chain mAb (clone MAR 1.8, BioXCell, 25 µg/day) over 3 consecutive days^54^. The last dose was administered 1 day before induction of I/R injury of the cremaster muscle or 1 h before intradermal (i.d.) MVA injection for skin analyses. The anti-Gr-1 mAb and anti-Ly6G mAb protocols reduced circulating neutrophil counts by >98% (n = 4-5 mice/group), as evaluated by flow cytometry in *Lyz2-eGFP-ki;Acta2-RFPcherry-Tg* mice 4 h after i.s. TNF or LPS administration or 24 h after MVA i.d. injection. Additionally, depletion efficiency was confirmed by analyzing neutrophil infiltration as indicated.

### Flow cytometry

Blood was collected from *Lyz2-eGFP-ki;Acta2-RFPcherry-Tg* mice in 5 mM EDTA in PBS and red blood cells were lysed using a commercial lysis buffer (BioLegend). Fc receptors were blocked by incubation with anti-CD16/CD32 antibodies (BioLegend). Samples were stained with anti-CD45-PerCP-Cy5.5 (Thermo Fisher Scientific) and anti-CD115-ATTO643 (BioLegend, labeled in-house) mAbs for 20 min at 4°C. Counting beads (BioLegend) were added, and neutrophil numbers/µL of blood were assessed using a FACSCelesta flow cytometer (BD) and FlowJo software (TreeStar). CD45^+^, eGFP^bright^, CD115^-^ cells were identified as neutrophils.

### GPIbα and CXCL7 inhibition and immunodepletion of GPVI

Blocking Fab fragments of the anti-GPIbα mAb (clone p0p/B, 2 µg/g body weight, produced in-house, blocking the vWF binding site^55^) or Fab control antibodies were i.v. injected 30 min before induction of cremasteric inflammation^56^. GPVI was depleted from platelets *in vivo* by i.p. injection of the anti-GPVI antibody, JAQ1 (50 µg/day, Emfret Analytics), or isotype-matched control mAbs on days 5 and 3 before induction of cremasteric inflammation. Where indicated, results from each control antibody treatment were similar and merged. Blocking anti-CXCL7 (clone 159742, 30 µg, R&D Systems) or corresponding isotype-matched control mAbs were injected i.s. together with TNF or at the beginning of the reperfusion phase in the I/R injury model for fixed tissue analyses. Circulating platelet and PMN counts after aforementioned antibody administration were determined at the end of the experiments using an automated hemocytometer. For IVM analysis, anti-CXCL7 mAb treatments are described in the corresponding section below.

### Infection of the ear skin

Mice were anesthetized with 3% isoflurane and i.d. injected with MVA (5 x 10^6^ IFU in 10 µL) or PBS alone^57,58^. 15 h or 24 h later, ear pinnae were harvested and fixed for immunostaining.

### Whole-mount immunofluorescence staining

Murine cremaster muscles or ear pinnae were fixed in 4% paraformaldehyde (Sigma-Aldrich) at 4°C for 1 h or 24 h, respectively. The ear pinnae were separated into dorsal and ventral sheets, and cartilage was mechanically removed. Tissues were blocked and permeabilized in PBS containing 3% bovine serum albumin (PanReac AppliChem) and 0.5% Triton X-100 (Carl Roth) for 4 h at room temperature. Tissues were stained with unlabeled or fluorophore-conjugated primary antibodies in 3% bovine serum albumin (BSA) in PBS overnight at 4°C. Where required, tissues were incubated with fluorescently labeled secondary antibodies in PBS containing 3% BSA for 3 h at room temperature. Antibody labeling with Alexa Fluor 488, 532, 555, or 647 was performed using commercial kits (Thermo Fisher Scientific) or by incubation with 10-fold excess of sulfo-Cy3 (Lumiprobe), ATTO 488 or ATTO 643 (ATTO-Tec) fluorochromes. Free dye was removed using 7 kDa Zeba-spin columns (Thermo Fisher Scientific), and labeling efficiency and antibody concentration were measured with a Nanodrop (Thermo Fisher Scientific).

### Confocal microscopy and image analysis

Immunostained whole-mounted cremaster muscles and ear sheets were imaged using inverted or upright TCS SP5 or SP8 confocal laser scanning microscopes (Leica Microsystems) equipped with argon ion, diode-pumped solid-state and helium-neon lasers. Excitation was performed at 405, 488, 561, and 633 nm. Images were acquired using water-dipping objectives (20x, 40x or 60x; NA 0.95-1.20) or oil immersion objectives (20x or 63x; NA 0.75-1.40). Serial z-stacks of postcapillary venules (diameter 20-45 µm) were acquired, and half-vessel images were reconstructed in 3D and analyzed using the Imaris v10.2.0 software (Bitplane, Oxford Instruments). Platelets, neutrophils, ECs, and pericytes were identified using antibodies against GPIX (clone p0p6, produced in-house), MRP14 (2B10, BD Biosciences), CD31 (polyclonal goat Ab, R&D Systems or *in vivo* stained by i.s. injection of clone 390, Thermo Fisher Scientific), or α-SMA (clone 1A4, Sigma-Aldrich), respectively.

Platelet numbers at different positions relative to the microvasculature were manually quantified within 300 x 150 x 45 µm fields of view from 8-10 images per mouse. Because reliable enumeration of individual platelets within the vessel lumen was hindered by platelet clusters, cumulative luminal platelet volume (representing luminal platelets tethered at the vascular segment) was measured instead, by generating iso-surfaces based on luminal regions immunostained for GPIX. Transmigrated neutrophils per 300 x 150 x 45 µm fields of view were quantified from the 8-12 images per mouse.

CXCL4 and CXCL7 levels within platelets were assessed using anti-CXCL4 (clone 140910, R&D Systems), anti-CXCL7 (goat polyclonal, R&D Systems), and corresponding fluorescently labeled secondary antibodies. MFI per unit area was quantified within GPIX^+^ platelet iso-surfaces. Chemokine MFIs were background-corrected by subtraction of the respective control signal obtained from samples incubated without primary antibodies. Chemokine MFIs and the proportion of chemokine-positive platelets were quantified from 4-10 images per mouse. Where indicated, sub-EC platelet MFIs were normalized to luminal platelet MFIs from the same mouse and expressed as percentages. All images for chemokine quantification were acquired using a 63x objective from 150 µm venular segments at a voxel size of 0.147 × 0.147 × 0.346 µm.

### Confocal IVM

Neutrophil and platelet dynamics and their interactions with the venular endothelial and pericyte layer were analyzed using confocal IVM in *Lyz2-eGFP-ki;Acta2-RFPcherry-Tg* mice (exhibiting eGFP^+^ neutrophils and RFP^+^ pericytes and SMCs^6,7^). Imaging parameters were configured to selectively detect eGFP^bright^ neutrophils^25^. EC junctions were *in vivo* labeled by i.s. injection of an Alexa Fluor 532-labeled non-blocking anti-CD31 mAb (clone 390, 4 µg, together with TNF^25^). Platelets were labeled by i.v. injections of an Alexa Fluor 647-conjugated non-blocking anti-GPIX mAb (clone p0p6, 10 µg, produced in-house^26^), 10 min before imaging. Calcium levels and dynamics in platelets were determined by IVM on *Pf4-Cre;PC::G5-tdT* mice (expressing tdTomato and the calcium sensor GCaMP5G in platelets and megakaryocytes), which received i.s. injection of an Alexa Fluor 647-anti-CD31 mAb (clone 390, 4 µg). 1-2 h after i.s. administration of TNF (300 ng) or 350 µL PBS alone, mice were placed on a heated, custom-designed microscope stage. Cremaster muscles were carefully exteriorized, incised, and pinned flat over the optical window of the stage and superfused with 37°C warm Tyrode’s solution (Sigma-Aldrich). In experiments involving platelet depletion and corresponding control treatments, any blood vessels disrupted by this procedure were carefully cauterized to prevent bleeding. For studying luminal neutrophil responses, blocking anti-CXCL7 mAbs (clone 159742, R&D Systems, 3 µg/g body weight) or isotype control mAbs were injected i.v. 10 min before TNF. For analysis of transendothelial migration and sub-EC neutrophil crawling, mAbs (30 μg) were injected i.s. 110 min after TNF administration^7^.

Postcapillary venules (20-45 µm diameter) were imaged for 20 min to 2 h using a TCS SP8 upright confocal laser scanning microscope with a 63x water-dipping objective (NA 0.9). Z stacks were acquired every 8 sec to quantify platelet calcium dynamics, 15 sec for analyzing platelet translocation across the EC layer, 30 sec to quantify luminal neutrophil responses, and 30 or 60 sec for analyzing sub-EC neutrophil and platelet responses. Image sizes were typically 100 × 65 × 27 µm for analyzing platelet dynamics and 300 × 130 × 35 µm for neutrophil tracking, resulting in voxel sizes of 0.05 × 0.05 × 0.8 μm or 0.29 × 0.29 × 0.69 μm, respectively, in x × y × z directions. Half vessels were reconstructed in Imaris to clearly visualize individual platelets and neutrophils and their interactions with the venular ECs and pericytes.

Luminal neutrophils were classified as adherent when stationary on the endothelium for ≥ 30 sec and as intraluminally crawling when exhibiting a displacement ≥ 2 cell diameters during the observation period and quantified from 4 vessels per mouse. Neutrophil transendothelial migration was defined as the complete passage of neutrophils through the endothelium in a luminal-to-abluminal direction and was quantified over 30 min. Sub-EC neutrophil and platelet dynamics were determined by manual tracking of individual cells in Imaris. The crawling straightness represents neutrophil displacement/track length. The fraction of sub-EC neutrophils breaching the pericyte layer was quantified within 15 minutes. Sub-EC platelets were defined as platelets with ≥50% of their body in the sub-EC compartment. Distances between sub-EC neutrophils and proximal platelets were determined using the spot-to-spot distance detection function and were measured between the centers of the cells. Platelet GCaMP5G (GFP) and tdTomato MFIs were quantified within platelet iso-surfaces, defined by tdTomato^+^ regions. Comparison between luminal and sub-EC platelet MFIs was performed at one time point per vessel and presented as MFI values for individual platelets.

### ELISA quantification of CXCL7

Cremaster muscles from mice treated with control or platelet-depleting antibodies (anti-GPIbα, R300) and i.s. stimulated with PBS or TNF (4 h) were snap frozen in liquid nitrogen. Tissues were homogenized in PBS containing 0.1% Triton X-100 (Carl Roth) and 1% protease inhibitor cocktail (Thermo Fisher Scientific) using a Bead Ruptor 12 (OMNI International), as described^59^. CXCL7 was quantified using a commercial ELISA kit (Raybiotech, sensitivity: 14 pg/mL). Total protein levels were determined using a commercial kit (Pierce™ Dilution-Free™ Rapid Gold BCA Protein Assay, Thermo Fisher Scientific). CXCL7 levels were expressed as ng/mg of total protein.

### Platelet counts, size, and surface receptor expression

Whole blood (50 µL) was collected into heparin (final concentration 20 U/mL) and diluted to 1 mL with Tyrode’s buffer without Ca^2+^. Aliquots (50 µL) were incubated with saturating concentrations of fluorophore-conjugated antibodies for 15 min at room temperature in the dark. Reactions were stopped by the addition of PBS, and samples were analyzed by flow cytometry. Platelet counts, forward scatter (as a surrogate of size), and MFI of surface receptors were determined.

### Platelet activation assays

For assessment of integrin αIIbβ3 activation and α-granule secretion, blood was collected and diluted as described above. To remove residual heparin, samples were washed twice in Tyrode’s buffer without Ca^2+^ (centrifugation 5 min at 800 *g*) and resuspended in Tyrode’s buffer supplemented with Ca^2+^. Washed samples were distributed into tubes containing fluorophore-conjugated antibodies against activated αIIbβ3 (clone JON/A-PE, Emfret Analytics) and P-selectin (clone WUG1.9, Emfret Analytics), together with the indicated agonists. Samples were incubated for 7 min at 37°C, followed by 7 min at room temperature in the dark. Reactions were stopped by the addition of PBS and analyzed by flow cytometry. Platelets were identified by forward and side scatter characteristics, and activation was quantified as MFI.

### Platelet adhesion and thrombus formation under flow

For analysis of platelet adhesion and thrombus formation on collagen, glass coverslips (24 × 60 mm) were coated overnight at 37°C with fibrillar type I collagen (200 µg/mL) and blocked with 1% BSA in PBS for at least 1 h at room temperature. Whole blood was collected into heparin (final concentration 20 U/mL) and diluted 2:1 with Tyrode’s buffer supplemented with Ca^2+^. Platelets were labeled with Alexa Fluor 647-conjugated anti-GPIX (0.3 µg/mL) for 5 min at 37°C before perfusion. Labeled blood was perfused over collagen-coated coverslips mounted in a transparent flow chamber (50 µm slit depth) at a shear rate of 1000 s^-1^ for 4 min using a pulse-free pump. Subsequently, Tyrode’s buffer supplemented with Ca^2+^ was perfused for 4 min to remove non-adherent cells. Thrombus formation was monitored by inverted microscopy. After washing, at least 8 randomly selected fields per coverslip were recorded and analyzed using ImageJ. Platelet surface coverage was quantified as percentage of total area covered, and thrombus volume was calculated as integrated fluorescence intensity (arbitrary units).

### Measurement of intracellular Ca^2+^ mobilization in platelets

Washed platelets were prepared as described above and adjusted to 2 × 10^8^ platelets/mL in Tyrode’s buffer supplemented with Ca^2+^. Platelet suspensions were loaded with Fura-2-AM (5 µM) in the presence of Pluronic F-127 (0.2 µg/mL) for 20 min at 37°C. Excess dye was removed by centrifugation (2 min, 800 *g*), and platelets were resuspended in HBSS containing 1 mM MgCl_2_ and 1 mM CaCl_2_.

Fluorescence measurements were performed under continuous stirring using alternating excitation at 340 and 380 nm, with emission recorded at 509 nm to determine baseline fluorescence. After baseline acquisition, agonists were added as indicated, and Ca^2+^ responses were recorded for 250 s. For calibration, 1% Triton X-100 was added to determine maximal fluorescence, followed by 0.5 M EGTA to determine minimal fluorescence, enabling calculation of absolute intracellular Ca^2+^ concentrations. Data were analyzed with Spectrum FL software and Microsoft Excel.

### Statistical analysis

Statistical analyses were performed using GraphPad Prism (version 10.4). The results are expressed as mean ± standard error of the mean (SEM) unless stated otherwise, and n numbers for each data set are provided in the figure legends. The Shapiro-Wilk test was used to determine the normal distribution of the data. For comparison between two groups, two-tailed Student’s t test with Welch’s correction was used for normally distributed data, and the two-tailed Mann-Whitney U test for non-normally distributed data. For comparisons of a single group against a normalized reference value, a two-tailed one-sample *t* test was used. For comparisons among more than two groups, one-way ANOVA followed by Dunnett’s or Tukey’s post hoc test or two-way ANOVA followed by Tukey post hoc test were carried out. Analyses were unpaired, unless stated otherwise in the figure legends. Statistical significance was accepted at p < 0.05.

