## Supplementary material for "Sub-endothelial platelet activation amplifies neutrophil transmigration within venular walls": Supplemetal Information

Supplemental Figures S1-S7

Table S1

Supplemental Video Legends (Videos S1-S5)

### Supplemental Figures

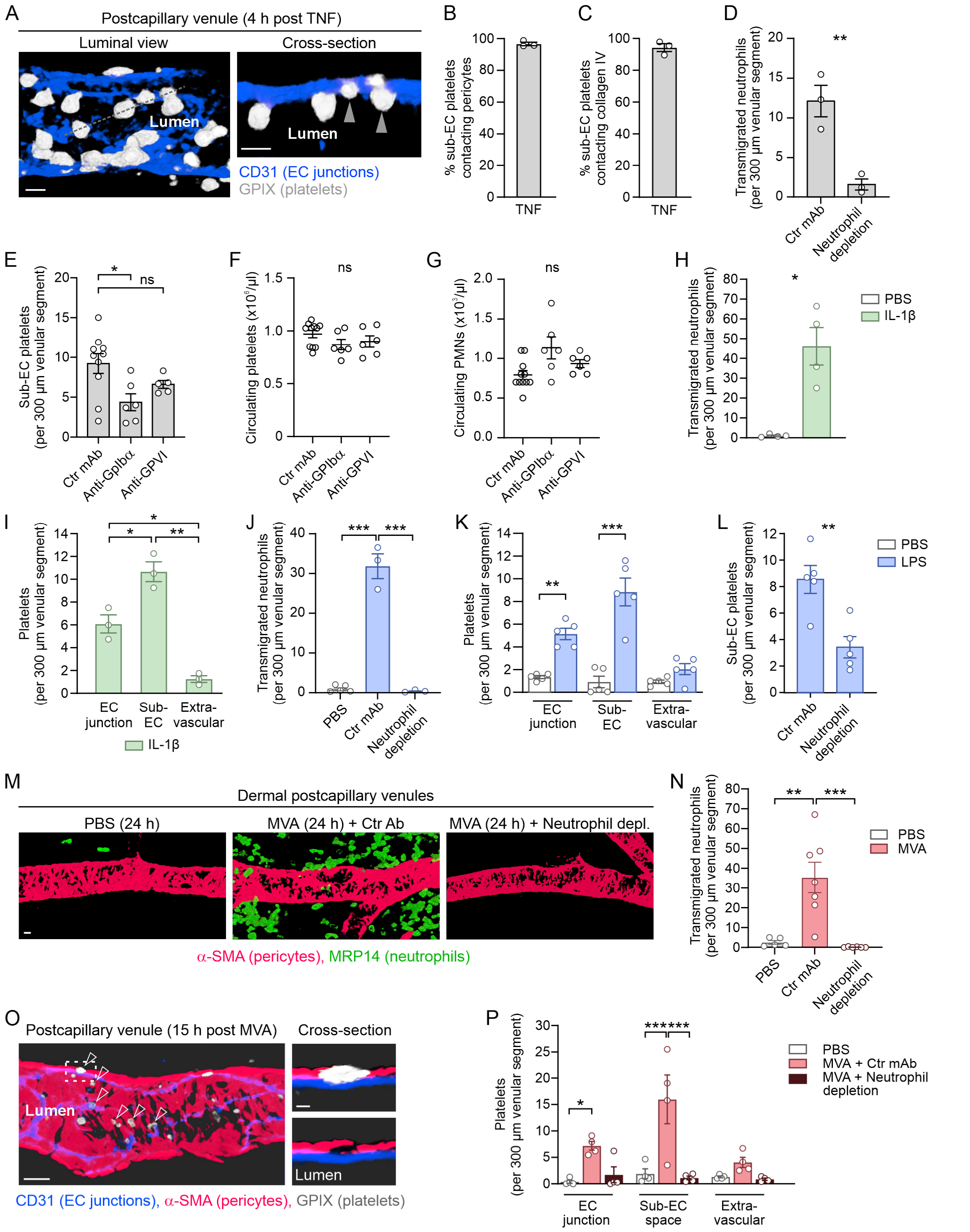


**Figure S1: Platelet localization in TNF-, IL-1β- and LPS-stimulated cremaster muscles, and the Modified Vaccinia Ankara virus (MVA)-infected skin. (A-D)** Cremaster muscles from wild-type (WT) mice were collected 4 h after PBS or TNF administration, immunostained for CD31 (endothelial cell (EC) junctions), GPIX (platelets), α-SMA (pericytes), MRP14 (neutrophils) and/or collagen IV, and analyzed by confocal microscopy. **(A)** Representative images of a postcapillary venule. Arrowheads indicate endothelial cell (EC) junctional platelets, defined as platelets within the vascular lumen protruding into EC junctions and quantified in Figure 1C. Scale bars, 5 µm. **(B, C)** Fractions of sub-EC platelets contacting **(B)** pericytes, and **(C)** collagen IV (n = 3 mice/group). **(D)** Number of transmigrated neutrophils in control or neutrophil-depleted mice treated with anti-Gr-1 mAbs (n = 3 mice/group). **(E-G)** Mice received control or anti-GPIbα Fab fragments (clone p0p/B; blocking the vWF-binding site) intravenously (i.v.), or control or anti-GPVI mAbs (clone JAQ1) intraperitoneally (i.p.) to immunodeplete GPVI from circulating platelets. **(E)** Number of sub-EC platelets (n = 5-10 mice/group), **(F)** circulating platelet counts, and **(G)** circulating PMN counts 4 h after TNF administration (n = 6-11 mice/group). **(H-I)** Cremaster muscles from control or neutrophil-depleted WT mice treated with anti-Gr-1 mAbs were analyzed by confocal microscopy 4 h after PBS, IL-1β, or LPS administration. **(H)** Numbers of transmigrated neutrophils and **(I)** platelets at distinct locations after PBS or IL-1β administration (n = 4 mice/group). **(J)** Number of transmigrated neutrophils (n = 3-5 mice/group), **(K)** platelets at distinct locations, and (**L**) number of sub-EC platelets after PBS or LPS administration (n = 5 mice/group). **(M-P)** Ear skin from control or neutrophil-depleted WT mice treated with anti-Ly6G mAbs was analyzed by confocal microscopy after intradermal PBS or MVA administration. **(M)** Representative images of a dermal venule. Scale bar, 10 µm. **(N)** Number of transmigrated neutrophils 24 h after PBS or MVA administration (n = 5-7 mice/group). **(O)** Representative dermal venule containing sub-EC platelets (arrowheads). Cross-sectional views of the boxed region are shown on the right. Scale bars, 10 µm (left) and 1 µm (right). **(P)** Platelets at distinct microvascular locations 15 h after PBS or MVA administration (n = 4 mice/group). Statistical significance was assessed using an unpaired two-tailed Student's *t* test with Welch's correction (D, H, L), one-way ANOVA followed by Dunnett's post hoc test (E-G and J), repeated-measures one-way ANOVA followed by Tukey's post hoc test (I), ordinary one-way ANOVA followed by Tukey's post hoc test (N), or two-way ANOVA followed by Tukey's post hoc test (K, P). Data are presented as mean ± SEM. *p < 0.05, **p < 0.01, ***p < 0.001; ns, not significant.

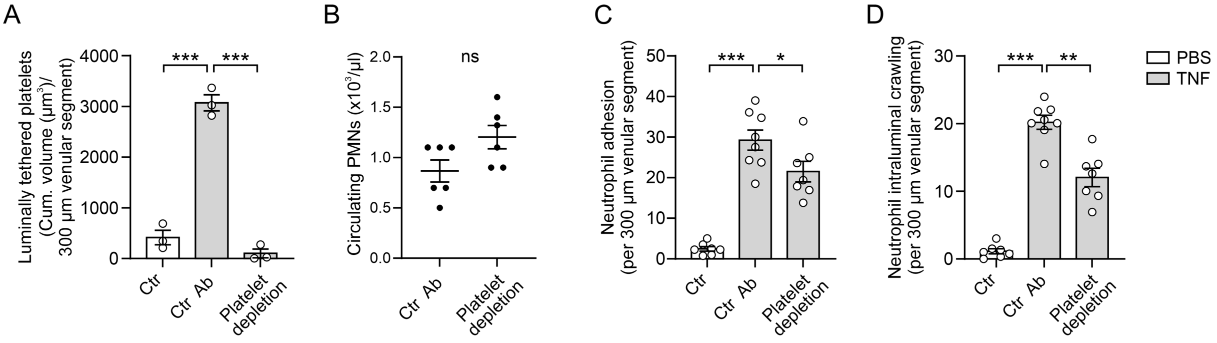


**Figure S2: Effects of platelet depletion on circulating platelet and PMN counts and on neutrophil adhesion and intraluminal crawling in TNF-stimulated cremaster muscles.**

**(A)** Cumulative volume of luminal platelets retained within cremasteric venular segments. WT mice received i.v. injections of control or platelet-depleting anti-GPIbα antibodies (R300), followed by intrascrotal (i.s.) administration of PBS or TNF. Cremaster muscles were collected after 4 h, immunostained, and analyzed by confocal microscopy (n = 3 mice/group). **(B)** Circulating PMN counts 4 h after TNF administration, determined using an automated hemocytometer (n = 6 mice/group). **(C, D)** *Lyz2-eGFP-ki;Acta2-RFPcherry-Tg* mice received control or platelet-depleting anti-GPIbα antibodies (R300), followed by i.s. PBS or TNF administration. Endothelial junctions and platelets were labeled *in vivo* with antibodies against CD31 and GPIX, respectively, and neutrophil dynamics in cremasteric venules were analyzed by confocal intravital microscopy (IVM) 2-4 h after TNF administration. **(C)** Neutrophil adhesion and **(D)** intraluminal crawling (n = 7-8 mice/group). Statistical significance was assessed using one-way ANOVA followed by Dunnett's post hoc test (A, C and D) or an unpaired two-tailed Student's *t* test with Welch's correction (B). Data are presented as mean ± SEM. $^{*}p < 0.05$, $^{**}p < 0.01$, $^{***}p < 0.001$; ns, not significant.

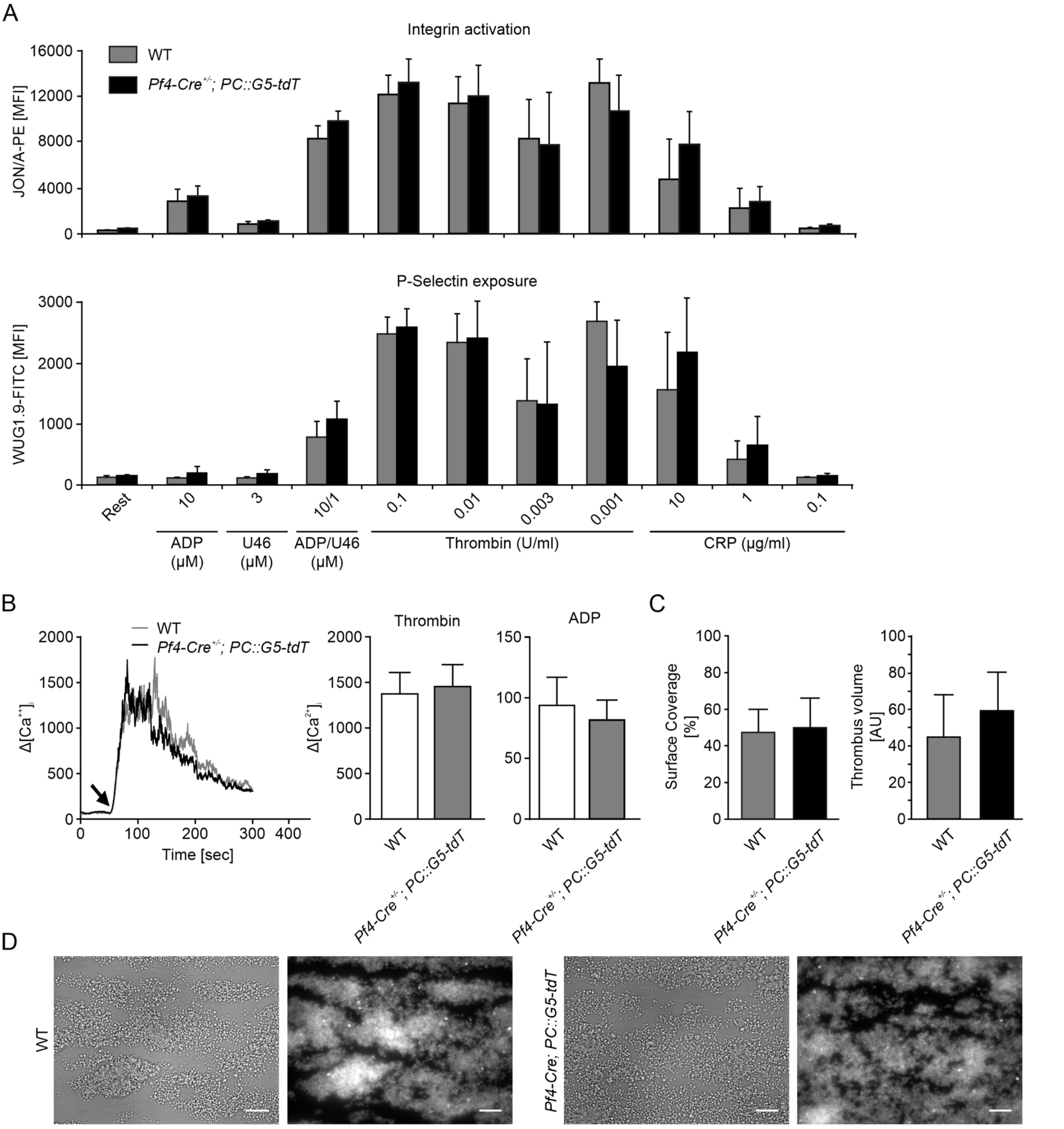


**Figure S3: Platelet-specific expression of GCaMP5G does not alter platelet function.**

**(A)** Flow-cytometric assessment of integrin αIIbβ3 activation (top) and α-granule-dependent P-selectin exposure (bottom) in platelets from WT and *Pf4-Cre;PC::G5-tdT* mice after stimulation with the indicated agonists. No significant differences were detected between genotypes. **(B)** Intracellular $Ca^{2+}$ mobilization in response to thrombin (0.01 U $ml^{-1}$; arrow indicates agonist addition) and ADP (5 µM) was comparable between genotypes. Left, representative $Ca^{2+}$ traces; right, maximal $Ca^{2+}$ responses (n = 4 mice/group). **(C, D)** Whole blood from WT and *Pf4-Cre;PC::G5-tdT* mice was labeled with Alexa Fluor 647-conjugated anti-GPIX antibodies (0.3 µg $ml^{-1}$) and perfused over collagen-coated coverslips at a shear rate of 1,000 $s^{-1}$. **(C)** Platelet surface coverage and thrombus volume (n = 5 mice/group). **(D)** Representative bright-field and fluorescence images. Scale bars, 20 µm. Statistical significance was assessed using a two-tailed Mann–Whitney U test. Data are presented as mean ± SD.

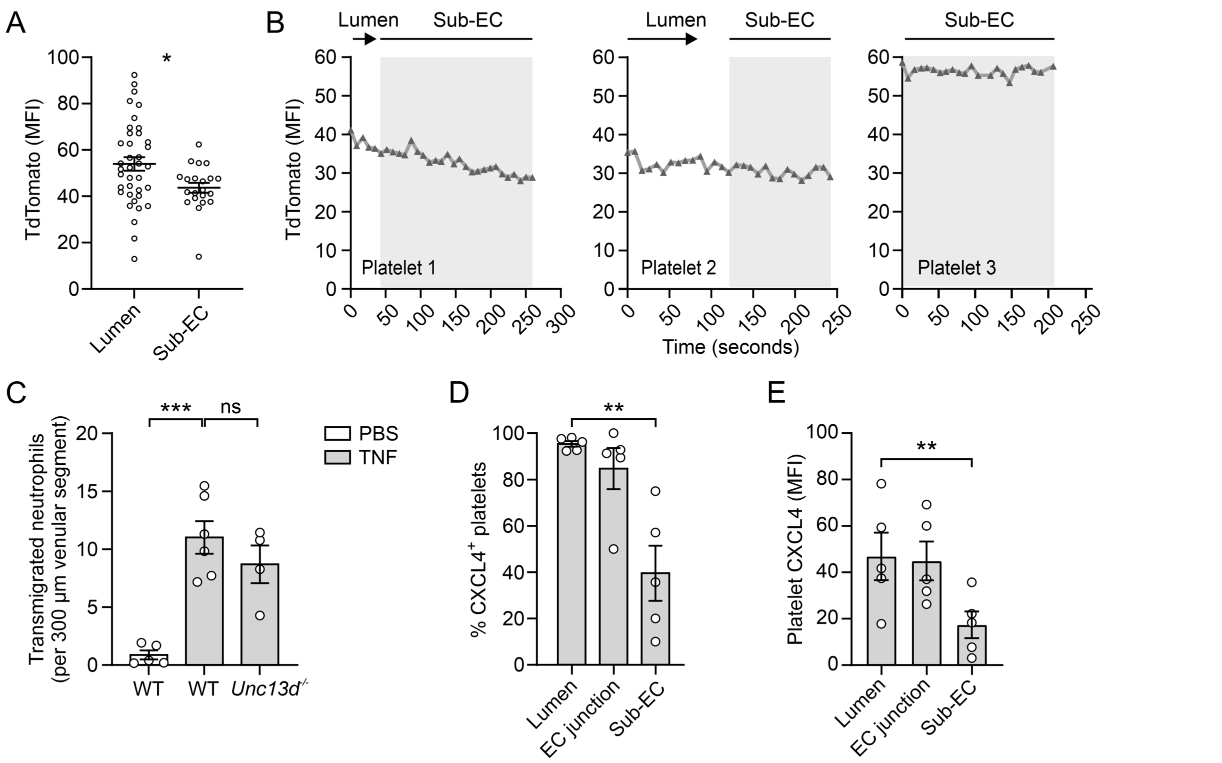


**Figure S4: Platelet tdTomato fluorescence, neutrophil extravasation in *Unc13d*$^{-/-}$ mice, and platelet CXCL4 levels in TNF-stimulated cremaster muscles.**

**(A, B)** Platelet tdTomato fluorescence was analyzed by confocal IVM in *Pf4-Cre;PC::G5-tdT* mice, which express tdTomato and the calcium indicator GCaMP5G in megakaryocytes and platelets, 2-4 h after i.s. TNF administration. Endothelial junctions were labeled *in vivo* with anti-CD31 mAbs. **(A)** TdTomato mean fluorescence intensities (MFIs) corresponding to the GCaMP5G measurements shown in Figure 4C, quantified in luminal and sub-EC platelets within TNF-stimulated cremasteric venules (n = 51 and 28 platelets pooled from 3 mice/group). MFIs were quantified at one time point per vessel and are shown for individual platelets. **(B)** Representative tdTomato fluorescence traces from individual platelets corresponding to Figure 4D. **(C)** Number of transmigrated neutrophils in cremaster muscles from WT and *Unc13d*$^{-/-}$ mice 4 h after PBS or TNF administration. *Unc13d*$^{-/-}$ platelets are deficient in dense-granule secretion (n = 4-6 mice/group). **(D, E)** Cremaster muscles from WT mice were collected 4 h after TNF administration, immunostained for CD31, GPIX and CXCL4, and analyzed by confocal microscopy. **(D)** Percentage of CXCL4$^{+}$ platelets and **(E)** CXCL4 MFI in platelets at distinct microvascular locations (n = 5 mice). Statistical significance was assessed using a two-tailed Mann–Whitney U test (A), one-way ANOVA followed by Dunnett's post hoc test (C), or repeated-measures one-way ANOVA followed by Dunnett's post hoc test (D, E). Data are presented as mean ± SEM. *$p < 0.05$, **$p < 0.01$, ***$p < 0.001$; ns, not significant.

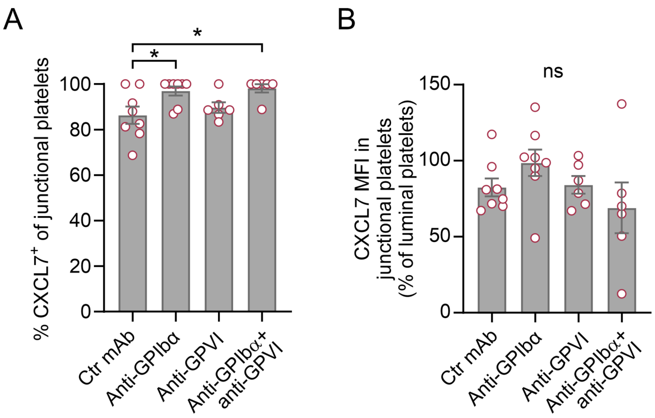


**Figure S5: CXCL7 levels in junctional platelets in TNF-stimulated cremaster muscles.**

WT mice received i.v. injections of control Fab fragments or Fab fragments of an anti-GPIbα mAb (clone p0p/B) 30 min before TNF administration, or were pretreated with control or anti-GPVI mAbs (clone JAQ1) 5 and 3 days before TNF administration to immunodeplete GPVI from circulating platelets. Cremaster muscles were collected 4 h after TNF administration, immunostained for CD31 (EC junctions), GPIX (platelets) and CXCL7, and analyzed by confocal microscopy. Results obtained with the corresponding control Fab and whole-IgG groups were similar and were therefore pooled. **(A)** Percentage of $CXCL7^+$ junctional platelets and **(B)** CXCL7 MFI in junctional platelets normalized to luminal platelets (=100%) in each mouse (n = 6-8 mice/group). Statistical significance was assessed using one-way ANOVA followed by Tukey’s post hoc test. Data are presented as mean ± SEM. *p < 0.05; ns, not significant.

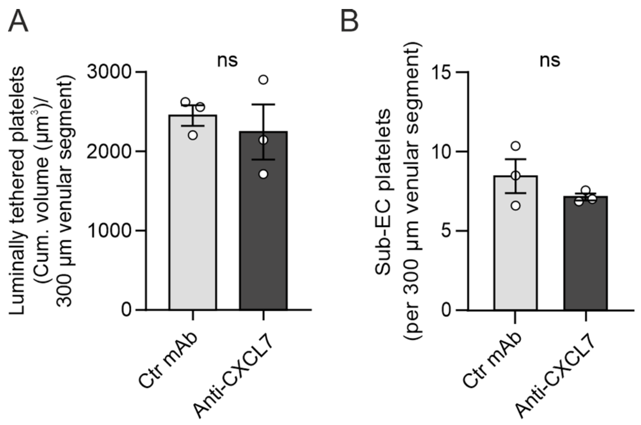


**Figure S6: CXCL7 blockade does not affect luminal platelet retention or sub-EC platelet accumulation in TNF-stimulated cremaster muscles.**

WT mice received i.s. injections of TNF together with control or blocking anti-CXCL7 mAbs (clone 159742). Cremaster muscles were collected 4 h later, immunostained for CD31, GPIX and α-SMA, and analyzed by confocal microscopy. **(A)** Cumulative volume of luminal platelets retained within venular segments and **(B)** number of sub-EC platelets (n = 3 mice/group). Statistical significance was assessed using an unpaired two-tailed Student's *t* test with Welch's correction. Data are presented as mean ± SEM; ns, not significant.

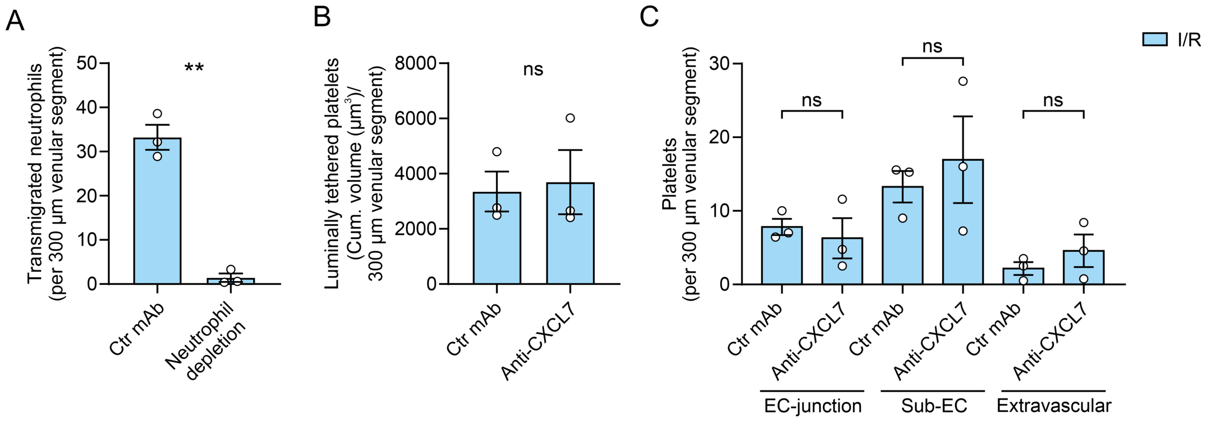


**Figure S7: Neutrophil depletion efficiency, and unchanged platelet localization after CXCL7 blockade in ischemia/reperfusion (I/R) injury-treated cremaster muscles.**

Cremaster muscles of WT mice were subjected to local I/R, stained for CD31, α-SMA, MRP14, GPIX, and analyzed by confocal microscopy. **(A)** Quantification of transmigrated neutrophils after control or anti-Ly6G mAb administration for neutrophil depletion. **(B)** Cumulative volume of luminal platelets, and **(C)** platelets at indicated locations after control or blocking anti-CXCL7 mAbs administration (n = 3 mice/group). Unpaired two-tailed Student's *t* test with Welch's correction (A and B), two-way ANOVA followed by Tukey's post hoc test (C). Means ± SEM, ns, not significant.

**Supplemental Table**

| | WT (MFI) | *Pf4-Cre; PC::G5-tdT* (MFI) | p-value |
|---|---|---|---|
| **GPV** | 5469 ± 142 | 5490 ± 237 | 0.8668 |
| **GPVI** | 1172 ± 186 | 1495 ± 390 | 0.1480 |
| **α2** | 1105 ± 65 | 1083 ± 27 | 0.5104 |
| **CD9** | 23690 ± 2579 | 24734 ± 546 | 0.4223 |
| **GPIb** | 7610 ± 374 | 7504 ± 145 | 0.5781 |
| **GPIX** | 9882 ± 122 | 10015 ± 274 | 0.3623 |
| **CLEC-2** | 3570 ± 190 | 3748 ± 274 | 0.2706 |
| **CD84** | 739 ± 247 | 1014 ± 255 | 0.1210 |
| **αIIbβ3** | 12369 ± 638 | 11561 ± 277 | 0.0446 |
| **α5** | 936 ± 112 | 1012 ± 16 | 0.2027 |
| **β3** | 4891 ± 295 | 4842 ± 203 | 0.7707 |

**Table S1. Surface expression of major platelet receptors is unaltered in *Pf4-Cre; PC::G5-tdT* mice.** Platelet surface receptor expression was analyzed by flow cytometry using FITC-conjugated antibodies. Mean fluorescence intensity (MFI) values are presented as mean ± SD. No significant differences were detected between WT and *Pf4-Cre;PC::G5-tdT* platelets (two-tailed Mann–Whitney U test; n = 5 mice/group).

**Supplemental Video Legends**

**Video S1. Sub-EC platelets in a TNF-stimulated cremaster muscle venule.** Three-dimensional confocal microscopy visualization of a venular wall from a WT mouse after i.s. TNF administration. Endothelial cell junctions (CD31, blue), pericytes (α-SMA, red), neutrophils (MRP14, green), and platelets (GPIX, grey) were visualized by immunostaining. Arrowheads indicate sub-EC platelets. Cross-sectional views show a platelet positioned between the endothelium and the pericyte layer.

**Video S2. Platelet entry into the sub-EC compartment following neutrophil transendothelial migration in a TNF-stimulated cremaster muscle venule.** Confocal IVM video of a cremasteric venule in the cremaster muscle of a *Lyz2-eGFP-ki;Acta2-RFPcherry-Tg* mouse after local TNF administration, showing eGFP$^{bright}$ neutrophils (green). Endothelial junctions (blue) and platelets (grey) were labeled *in vivo* with Alexa Fluor 532-conjugated anti-CD31 and Alexa Fluor 647-conjugated anti-GPIX mAbs, respectively. The video shows a neutrophil undergoing transendothelial (trans-EC) migration, followed by a platelet adhering to the luminal endothelial surface and subsequently traversing the endothelium through the same junctional site. For clarity, a representative neutrophil and platelet are highlighted by isosurface rendering in Imaris. Still images from this video are shown in Figure 1G.

**Video S3. Neutrophil sub-EC crawling toward sub-EC platelets in a TNF-stimulated cremasteric venule.** Confocal IVM movie of a TNF-stimulated cremasteric venule in a *Lyz2-eGFP-ki;Acta2-RFPcherry-Tg* mouse, showing eGFP$^{bright}$ neutrophils (green) and RFP$^{+}$ pericytes (red). Endothelial junctions (blue) and platelets (grey) were labeled *in vivo* with Alexa Fluor 532-conjugated anti-CD31 and Alexa Fluor 647-conjugated anti-GPIX mAbs, respectively. The video follows a neutrophil undergoing transendothelial migration into the sub-EC compartment, crawling toward sub-EC platelets, and subsequently exiting the venular wall in their immediate vicinity. For clarity, the tracked neutrophil and sub-EC platelets are shown following isosurface rendering in Imaris. Two sequences of the same event are presented. Still images are shown in Figure 2B.

**Video S4 (related to Figure 4). GCaMP5G-reported calcium signals in a TNF-stimulated cremaster muscle venule.** Confocal IVM of a cremasteric venule in a *Pf4-Cre;PC::G5-tdT* mouse after TNF administration showing tdTomato$^{+}$ platelets (grey) and intracellular $Ca^{2+}$ signals reported by GCaMP5G (pseudocolor map of pixel-wise fluorescence intensity). Endothelial junctions (blue) were labeled *in vivo* with an Alexa Fluor 647-conjugated anti-CD31 mAb. Circles indicate sub-EC platelets. Still images are shown in Figure 4B.

**Video S5 (related to Figure 4). GCaMP5G-reported calcium dynamics during platelet entry into the sub-EC compartment.** Confocal IVM of a cremasteric venule in a *Pf4-Cre;PC::G5-tdT* mouse after i.s. TNF administration, showing tdTomato$^{+}$ (grey), and intracellular $Ca^{2+}$ signals reported by GCaMP5G (pseudocolor map of pixel-wise fluorescence intensity). Endothelial junctions (blue) were labeled *in vivo* with an Alexa Fluor 647-conjugated anti-CD31 mAb. The video shows a platelet translocating from the lumen into the sub-EC compartment. For clarity, the platelet is shown following isosurface rendering in Imaris. Two sequences of the same event are presented: the first shows platelet entry into the sub-EC compartment, and the second shows the corresponding $Ca^{2+}$ dynamics, with the platelet indicated by a circle. Corresponding GCaMP5G and tdTomato fluorescence traces are shown in Figure 4D (platelet 1).
